# Understanding prefrontal cortex functions by decoding its molecular, cellular and circuit organization

**DOI:** 10.1101/2022.12.29.522242

**Authors:** Aritra Bhattacherjee, Chao Zhang, Brianna Watson, Mohamed Nadhir Djekidel, Jeffrey R. Moffitt, Yi Zhang

**Affiliations:** Howard Hughes Medical Institute, Boston Children’s Hospital, Boston, Massachusetts 02115, USA; Program in Cellular and Molecular Medicine, Boston Children’s Hospital, Boston, Massachusetts 02115, USA; Division of Hematology/Oncology, Department of Pediatrics, Boston Children’s Hospital, Boston, Massachusetts 02115, USA; Department of Microbiology, Harvard Medical School, Boston, Massachusetts 02115, USA; Center for Applied Bioinformatics, St. Jude Children’s Research Hospital, 262 Danny Thomas Place, Memphis, TN, 38105, USA; Department of Genetics, Harvard Medical School, Boston, Massachusetts 02115, USA; Harvard Stem Cell Institute, WAB-149G, 200 Longwood Avenue, Boston, Massachusetts 02115, USA

**Keywords:** Mouse prefrontal cortex, spatial transcriptomics, MERFISH, neuron cluster projection, pain

## Abstract

The prefrontal cortex (PFC) is functionally one of the most complex regions of mammalian brain. Unlike other cortical areas that process single sensory modalities (like vision, touch, smell, etc.), the PFC integrates information across brain regions to regulate diverse functions ranging from cognition, emotion, executive action to even pain sensitivity. However, it is unclear how such diverse functions are organized at the cellular and circuit levels within the anatomical modules of the PFC. Here we employed spatially resolved single-cell transcriptome profiling to decode PFC’s organizational heterogeneity. The results revealed that PFC has very distinct cell type composition relative to all neighboring cortical areas. Interestingly, PFC also adopts specialized transcriptional features, different from all neighbors, with differentially expressed genes regulating neuronal excitability. The projections to major subcortical targets of PFC emerge from combinations of neuron subclusters determined in a target-intrinsic fashion. These cellular and molecular features further segregated within subregions of PFC, alluding to the subregion-specific specialization of several PFC functions. Finally, using these unique cellular, molecular and projection properties, we identified distinct cell types and circuits in PFC that engage in pathogenesis of chronic pain. Collectively, we not only present a comprehensive organizational map of the PFC, critical for supporting its diverse functions, but also reveal the cluster and circuit identity of a pathway underlying chronic pain, a rapidly escalating healthcare challenge limited by molecular understanding of maladaptive PFC circuits.

**Major points:**

- PFC adopts unique cellular composition, distinct from other cortical areas
- Selective transcriptomic features emerge in PFC to support its divergent functional portfolio
- Subcortical projections of PFC assume target-intrinsic specification for innervating clusters
- A molecularly defined L5 projection neuron cluster (to PAG) potentially mediates chronic pain pathogenesis

## Introduction

The prefrontal cortex (PFC) is a major region of the mammalian brain that has evolved to perform highly complex behavioral functions. It plays important roles in cognition, emotion and executive function. Unlike somatosensory, visual, auditory, motor or other cortices, which are unimodal (process single modalities like touch, vision, hearing, movement etc.), the PFC engages in complex executive tasks that dynamically coordinate cognition, attention, learning, memory, judgement, etc. to direct the action of an organism^1,2^. As such, dysfunctions of the PFC are associated with many cognitive and neuropsychiatric disorders^3,4^.

In addition to regulating intellectual and emotional behaviors, PFC is even involved in modulating pain sensitivity as well as the negative affect of pain^5,6^. Increasing evidence indicates that disruption of this regulation is associated with the development of chronic pain, a rapidly increasing healthcare challenge that affects about 20% of the US population, exceeding cost burden of diabetes or heart disease^7,8^. Chronic pain has been associated with PFC hypoactivity, and transcranial stimulation of the PFC can induce pain relief^9–15^. Although projections from PFC to brainstem has been historically described in descending inhibition of pain^5,16,17^, the underlying molecular mechanism is poorly characterized. Besides, PFC interacts with many downstream targets including the amygdala, nucleus accumbens and thalamus - the major components of the central pain matrix, critical for the sensory or affective symptoms of chronic pain^6,18^. As such, PFC plays an important role in pain “chronification”^5,17^.

Thus, a central question is how does PFC organize and manage such diverse functions: from cognitive processes to autonomic pain modulation? To address this question, we and others have previously performed single-cell RNA-seq (scRNA-seq) to decode the cellular heterogeneity of the PFC^2,19^, which revealed a myriad of cell types comprising PFC. However, those studies lacked information about the spatial organization and interaction of the diverse cell types, which are major determinants of the functional diversity of the PFC. A relatively homogeneous histology, with a laminar organization, is the most striking feature of the mammalian cerebral cortex^20–22^. Yet, distinct regions of cortex perform highly specialized functions, including vision, locomotion, and somatosensation, etc. This regional specialization of functions, despite apparent homogeneity, must be due to distinct features at multiple levels including - molecular composition (transcriptome), circuit organization (connectome) and anatomical (spatial) organization of cell subtypes within each cortical area. Decoding such organizational logic is critical not only for mechanistic understanding of cortical function, but also for developing drugs to selectively target neurological disorders of cortical origin, such that drugs directed for either cognitive (frontal cortex) or hearing (auditory cortex) defect, do not disrupt visual or motor function.

Approaching such questions has been historically limited by technological barriers, despite extensive scRNAseq profiling across brain, including cortex^23–25^. However, with recent advances in spatial transcriptomics techniques, such questions can now be addressed. Using multiplexed error-robust fluorescence in situ hybridization (MERFISH), an image-based method for spatially resolved single cell transcriptomics^26,27^, here we vividly decode the spatial organization of the PFC and its various subregions. Our results demonstrate distinct cellular composition of the PFC relative to its adjoining cortical areas. PFC adopts unique molecular features to suit its specific electrophysiological properties different from its adjacent motor cortex. We map molecular identities (and layer localization) of projection neurons to major subcortical targets. Finally, based on projection, transcription and activity marks, we reveal the molecular identity of PFC clusters most significantly affected in chronic pain.

## Results

### MERFISH reveals molecular diversity and location of cell types in the frontal cortex

To understand the diversity of cell types and determine their spatial organization within the PFC, we performed MERFISH^26 27–29^, the imaging-based method for single cell transcriptomics that uses combinatorial labeling of RNA species with error-robust barcoding which are read through iterative rounds of single-molecule FISH. MERFISH detects the precise location of each RNA molecule to ultimately reveal the spatial organization of diverse cell types within anatomically defined tissue regions (Fig. S1a)^29 28^. We constructed a MERFISH library to interrogate 416 genes consisting of cell-type markers and functionally important genes including-ion channels, neuropeptides, G-protein coupled receptors and a panel of neuronal activity regulated genes (Supplementary Table 1). We collected brain samples from three different mice and prepared rostral to caudal coronal slices covering +2.5 to +1.3 from Bregma to broadly image the frontal cortex. Using established analysis pipelines^29^, imaged RNA species were detected, decoded and assigned to individual cells by segmentation based on poly(A) and nuclear (DAPI) staining (Fig. S1a). Overall, we obtained 487,224 high-quality cells in the frontal cortical region from three independent biological replicates with high consistency (Fig. S1b). Expression of individual genes showed good correlation with that of the bulk RNA-seq of the PFC (Fig. S1c).

After unsupervised clustering, we identified the major cell types including excitatory neurons, inhibitory neurons, and non-neuronal cells that include oligodendrocytes, oligodendrocyte precursors (OPC), microglia, endothelia, astrocytes and vascular leptomeningeal cells (VLMC) (Fig. 1a). Within the excitatory neurons, the major subgroups clustered together, as described by the commonly used nomenclature^23^: the intra-telencephalic (IT) populations of different layers, the extra-telencephalic (ET) neurons, the near projecting (NP) and the cortico-thalamic (CT) populations (Fig. 1a). Within the inhibitory neurons, populations from the medial ganglionic eminence (*Pvalb* and *Sst*) and the caudal ganglionic eminence (*Vip*, *Sncg* and *Lamp5*) clustered distinctly (Fig. 1a).

**Fig. 1:**
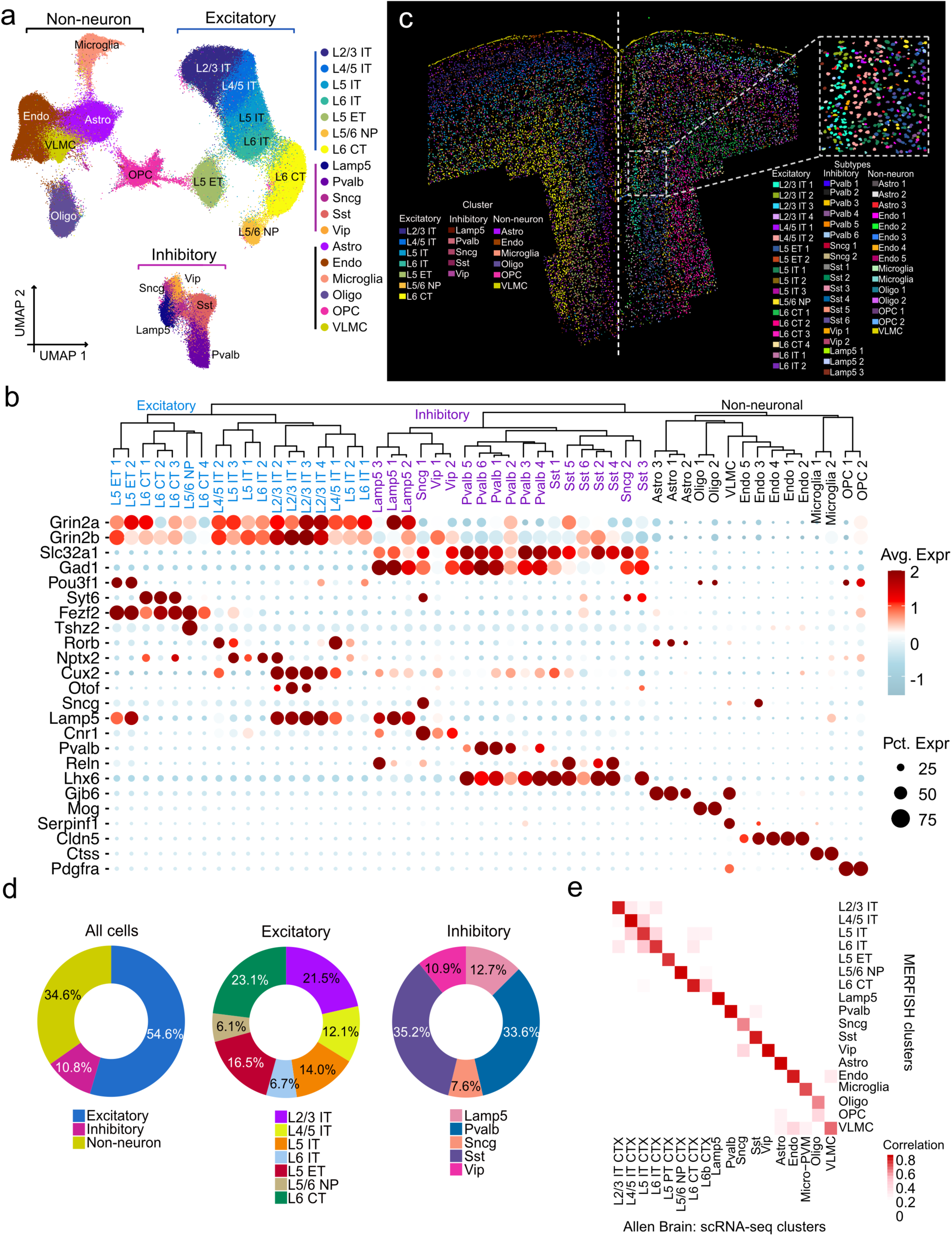
MERFISH reveals the molecularly diverse cell types and subtypes comprising the PFC. **a**, UMAP visualization of all cells identified by MERFISH. Cells are colored coded by their identities. **b**, Dendrogram showing the hierarchical relationship among all molecular defined cell subtypes. The expression of marker genes is shown below. The color represents the average expression, and dot size indicates the percentage of cells expressing each gene. **c**, Spatial map of all cell subtypes in a represented coronal slice. An enlarged view of a zoom-in region is shown in the top-right. **d**, Pie charts showing the cell proportions of the major cell types (left), excitatory neurons (middle) and inhibitory neurons (right) in PFC. **e**, Heatmap showing the gene-expression correlation between cell types and subtypes defined by MERFISH and scRNA-seq. scRNA-seq data are downloaded from Allen brain atlas, and only cells from PFC are used.

The major cell types were further clustered into 52 hierarchically organized cell subtypes, including 18 excitatory neuron subtypes, 19 inhibitory neuron subtypes, and 15 non-neuron subtypes (Fig. 1b). Among the excitatory IT neurons, four subtypes were detected in L2/3 (L2/3 IT 1 to 4), two subtypes in L4/5, three subtypes in L5, and two subtypes in L6. Additionally, the L5 ET split in two subtypes, the L6 CT into four subtypes and L5/6 NP formed a single cluster. Among the inhibitory neurons, the Pvalb and Sst each split into six subtypes, the Lamp5 into three subtypes, the Vip and Sncg into two subtypes each (Fig. 1b). Among the non-neurons, the endothelial cells formed five subtypes, Endo1-5, while astrocytes formed three subtypes, oligodendrocytes and OPCs each formed two subtypes.

Projecting these clusters in space (based on MERFISH coordinates) revealed the anatomical layout of the coronal section and depicted precise localization of every single cell (Fig. 1c; inset – magnified view showing individual cells). We found that molecularly similar excitatory neurons are localized together in space to form distinct layers, from which a laminar histology, characteristic of cerebral cortex, emerged (L2/3 IT to L6 CT: outside inwards) (Fig. 1c, left half). Within each layer, the subtypes are further organized in strata (e.g., L2/3 IT 1 to L2/3 IT 4) (Fig. 1c, right half: distribution of subtypes). Inhibitory neurons are broadly distributed and do not form specific layers, although some subtypes appear to be enriched within certain layers or subregions (Fig. 1c). Non-neuronal cells are also broadly distributed, except for enrichment of oligodendrocytes near the fiber tracts (e.g., corpus callosum) and the VLMC in the outermost layer of brain (Fig. 1c, yellow). As evidence of accurate localization, we mapped the RNA location of a few well-characterized genes that are known to express only in specific cortical layers. We found *Otof*, *Cux2* and *Fezf2* mRNAs are respectively localized to L2, L2/3 and L5 on the MERFISH slice, consistent with the ISH Images from Allen Brain Institute (Fig. S1d)

Together, excitatory neurons comprise the largest population in PFC, followed by non-neuronal cells combined and then the inhibitory neurons (Fig. 1d, left). Within excitatory, the IT neurons are the largest subgroup, followed by the ET, NP and CT of deeper layers, respectively (Fig. 1d, middle). Within the inhibitory, Sst and Pvalb neurons are most abundant followed by the Lamp5, Scng and Vip (Fig. 1d, right).

To further evaluate our detection accuracy, we first performed an integrated analysis of the MERFISH data with scRNA-seq data of the PFC from the Allen Institute^23^. All the major subtypes showed strong correlation between the two datasets (Fig. 1e). Similar integrated analysis comparing the MERFISH data with our own scRNA-seq of PFC^19^ revealed strong correspondence even at the subtype levels (Fig. S2a-e, see Method). In fact, MERFISH could classify some of the scRNA-seq clusters at a finer resolution to reveal distinct subclusters (Fig. S2d, e). This point is particularly true for the inhibitory neurons (e.g. Inh 1, 2 and 7 of scRNA-seq), possibly due to their higher rate of detection in MERFISH (Fig. S2f).

Collectively, our results indicate that we have faithfully detected all known cell subtypes and their locations within the mouse frontal cortex, which enables us to analyze their spatial organization along all 3D axes within this region.

### Marked heterogeneity in spatial distribution of neuron subtypes along AP and DV axes in PFC

To understand spatial organization of the different neuron subtypes within the anatomically defined PFC region, we aligned our profiled frontal cortex sections with the Allen Mouse Brain Common Coordinate Framework (CCFv3)^30^, a reference created for the mouse brain based on serial two photon tomography images of the 1675 C57Bl6/J mice (Fig. S3a), which outlines the PFC-boundaries within each section.

Mapping the MERFISH clusters onto the sequential antero-posterior sections revealed the order of cellular organization in 3D throughout the frontal cortex (Fig. 2a). Heterogeneous distribution of several neuron subtypes along the antero-posterior (A-P) and dorsal-ventral (D-V) axes was visually evident. Analysis along the AP axis revealed that L2/3 IT and L4/5 IT neuron subtypes are most enriched in the anterior-most part of the frontal cortex, where all types of L5 and L6 neurons are generally low (Fig. 2b). This density gradient follows a reverse order in posterior direction where deep layer neurons like L5 ET 1 or L6 CT 1-3 are gradually enriched (Fig. 2b). Detailed mapping of various neuron subtypes on the serial brain sections clearly revealed the uneven distribution along the A-P axis (Fig. 2c, S3b). In contrast, some subtypes such as L5/6 NP are modularly distributed and few others (e.g., L5 IT 2 or L6 IT 1) are sparse, but uniform throughout the A-P axis (Fig. 2b). IT neurons generally project to shorter distances within the telencephalon or cortex, while non-IT neurons predominantly project long distances outside telencephalon. This distribution likely favors the anterior bias of IT cells and the posterior bias of non-IT neurons closer to the subcortical region and the major fiber tracts.

**Fig. 2:**
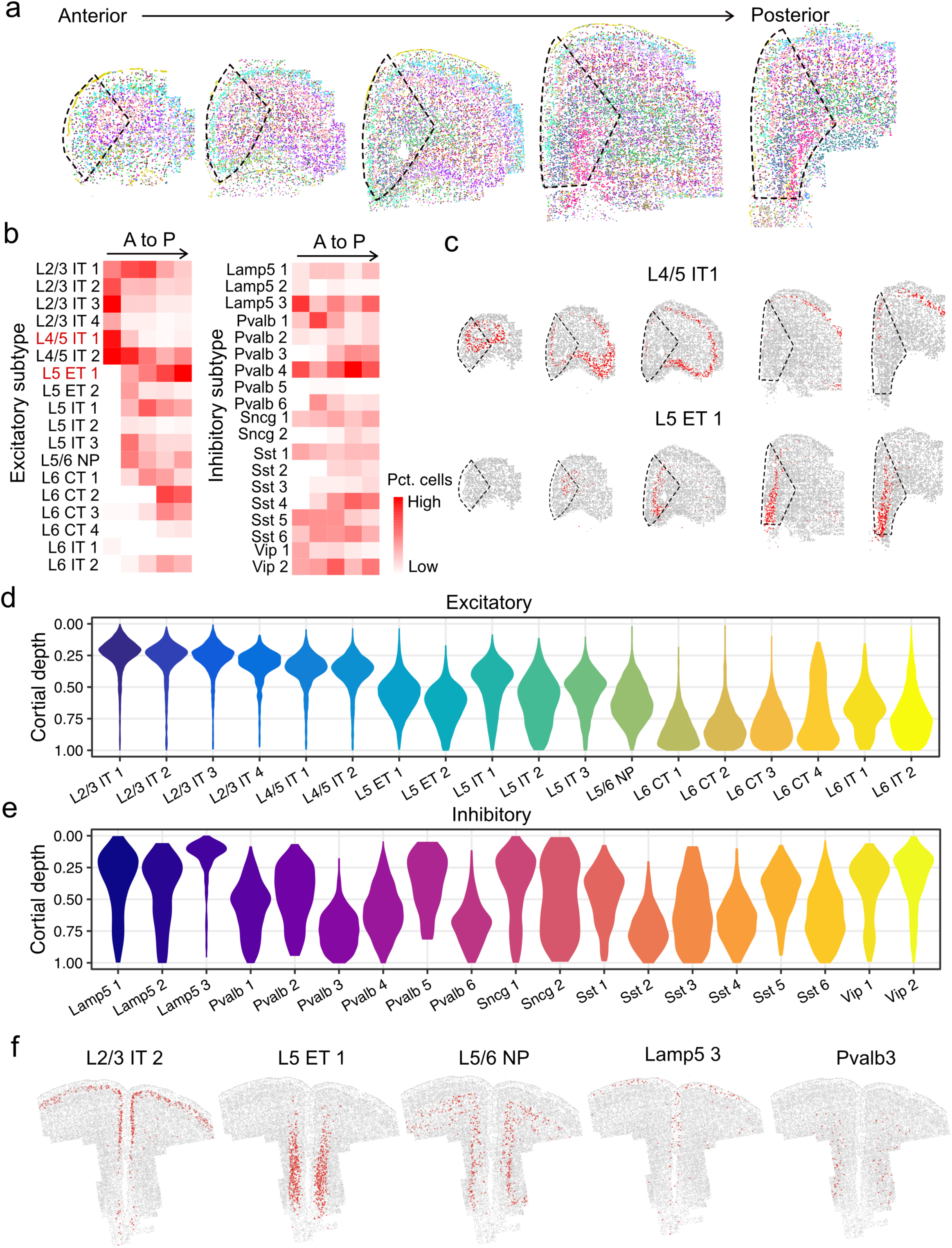
Spatial organization of different neuron subtypes in PFC. **a**, Coronal MERFISH slices showing the spatial organization of neuron subtypes from anterior to posterior in PFC and adjacent regions. The dotted lines indicate the PFC region. The color scheme is the same as in Fig. 1c. **b**, Heatmap showing the proportions of neuron subtypes within PFC from anterior to posterior (A to P) sections in excitatory (left) and inhibitory (right) neurons. **c**, Spatial organization of L4/5 IT 1 and L5 ET 1 from anterior to posterior sections. **d**, **e**, Violin plots showing the cortical depth distributions of excitatory neuron subtypes (**d**) and inhibitory neuron subtypes (**e**) in PFC. The maximum cortical depth is normalized to 1. **f**, Spatial location of five representative neuron subtypes (excitatory neuron subtypes: L2/3 IT 2, L5 ET 1 L5/6 NP; and inhibitory neuron subtypes: Lamp5 3, Pvalb 3) on a coronal slice. Red dots mark the indicated cell types and gray dots mark the other cells.

There is also strong distribution heterogeneity among the inhibitory neurons, but it follows a pattern of regional enrichment instead of gradual transitions along the A-P axis (Fig. 2b). For some subtypes, such as Lamp5 3, Pvalb 4 and Vip 2, the fluctuation in density along the A-P axis is very prominent (Fig. 2b). Neighborhoods with high density of distinct interneuron subtypes may indicate regulatory hotspots and/or focal points for specific subcortical projections circuits. Such unique organization reaffirms the principle that function begets structure in biological systems. It is only through spatial transcriptomics that such information can be accurately revealed.

Another readily recognizable feature from the coronal slices is the laminar organization of various excitatory neurons along the DV axis, within each representative section (Fig. 2a). Computation of physical depth inward from the cortical surface revealed that IT neurons locate more superficially within each layer. The L2/3 IT (and L4/5 IT) subtypes are most superficial and closer to the surface of the brain (Fig. 2d). Similarly, in L5, most IT neurons (L5 IT 1, L5 IT 3) are superficial to the other populations of the layer (L5 ET 1, L5 ET 2) (Fig. 2d). Within Layer 6, although L6 IT 1 is superficial, L6 IT 2 mingles with the deepest CT subtypes (Fig. 2d). Plotting each population individually onto a representative coronal section, a highly specific spatial localization of each neuron subtypes in layers inwards from the surface is clearly resolved (Fig. 2f, S4a). This layered organization is precisely achieved during developmental migration of cortical neurons when the migrating wave of each cell type is regulated by cues originating from their final homing site/layer^31^. As such, the types and density of neurons can be influenced by local signals to form circuits or hotspots characteristic of distinct subregions. How such anatomic assemblies locally emerge, remains a matter of further study.

The DV organization of GABAergic interneurons was even more interesting. Although, inhibitory neurons, unlike the excitatory, are not organized in layers, most subtypes appear to be enriched within specific excitatory layers or subregions (Fig. 2e). Broadly, the Lamp5 (Lamp5 1 to 3) and Vip (Vip 1 and 2) neurons along with Sncg 1 are more enriched in superficial layers. Lamp5 3, for example, is restricted only to the superficial layer (Fig. 2e, f). However, Sncg 2 is broadly distributed along the entire depth (Fig. 2e, S4b). This appears to be different from neighboring motor cortex, as per recent reports^32^, where all subtypes of *Sncg* neurons are present only in superficial layers. Additionally, motor cortex also has some subtypes of *Vip* neurons in deeper layers, which was not detected in PFC. However, the most interesting observation is that specific molecular subtypes of Pvalb and Sst neurons are differentially enriched in various layers along the cortical depth (Fig. 2e). For example, while Pvalb 5 and Pvalb 2 have higher density towards the superficial layers, Pvalb 3 and Pvalb 6 are enriched in the very deep layers, and Pvalb 1 and Pvalb 4 are maximally enriched in the intermediate region (Fig. 2e, 2f, S4b). Likewise, Sst 1 and Sst 5 are more superficially enriched, and the remaining are distributed in the intermediate to deep layers (Fig. 2e, S4b). Pvalb neurons can regulate excitatory pyramidal neuron firing through feedforward inhibition delivered directly onto the somatic compartment, while Sst neurons targets distal dendrites of excitatory neurons to impose feedback inhibition^33,34^. Although these interactions are indispensable to calibrate cortical excitatory output^35^, it is striking that inhibitory neurons diversified distinct molecular subtypes to adapt to the molecular diversity of excitatory pyramidal neurons in each cortical layer.

Most non-neuronal subtypes displayed a more broad and dispersed distribution (Fig. S4c), with few exceptions. The vascular leptomeningeal cells (VLMC), for example, line the outermost surface along the cortex. The Oligo 1 and Oligo 2 are enriched near the regions of origin of the white matter tracts (Fig. S3c). The Astro 2 had significant presence in L1 and somewhat greater enrichment in the medial prefrontal region (Fig. S4c).

### Distinct neuron subtypes are uniquely enriched in PFC

PFC is very distinct in function and connectivity compared to the adjacent cortices. We asked whether this functional and connectivity distinction is associated with its specialized cell composition. To this end, we identified the PFC boundary in each section by aligning with CCFv3 (Fig. S3a). By projecting the cells identified from the alignment as ‘in’-PFC onto the combined UMAP of frontal cortex (Fig. 3a), we found that some subtypes of excitatory neurons are selectively biased ‘in’, and some others ‘out’ of the defined PFC region (Fig. 3a), indicating different cellular composition in PFC and the adjacent areas. Relative population enrichment calculation showed that L2/3 IT 1, L5 ET 1 and L5 IT 1 are about 8 folds enriched within the PFC, whereas L6 CT 2 and L6 CT 3 are enriched by more than 2 folds (Fig. 3b). In contrast, L2/3 IT 4, L4/5 IT 1 or L6 IT 1 are markedly depleted (4-8 folds) in the PFC (Fig. 3b). When mapped onto the representative coronal section, the enriched, depleted and unbiased populations were clearly visible with respect to the boundaries of the PFC (Fig. 3c). Inhibitory neurons, although less abundant, exhibit clear subtype selectivity across all the major types in PFC (Fig. 3b). Switching of Pvalb subtypes (~2 fold enriched in Pvalb 3 and 4, and depleted in Pvalb 1, 2, and 6), depletion of Sncg 2 and enrichment of Sst 4 and 6, are the most prominent features (Fig. 3b, S5a). Also notably, Lamp5 3, the most superficially located interneuron (L1) is the only enriched Lamp5 neuron in PFC (Fig. 3b). The relative proportions of specific IT, ET and CT subtypes are intimately tied to the projections of a cortical area (inside and outside the telencephalon). The selection of specific interneurons determines the precise excitatory-inhibitory balance in the input/output circuits of the projections. In combination, these circuit motifs likely serve as a blueprint for the specialized functions of a cortical area, and PFC is clearly organized into a highly selective assembly in this regard.

**Fig. 3:**
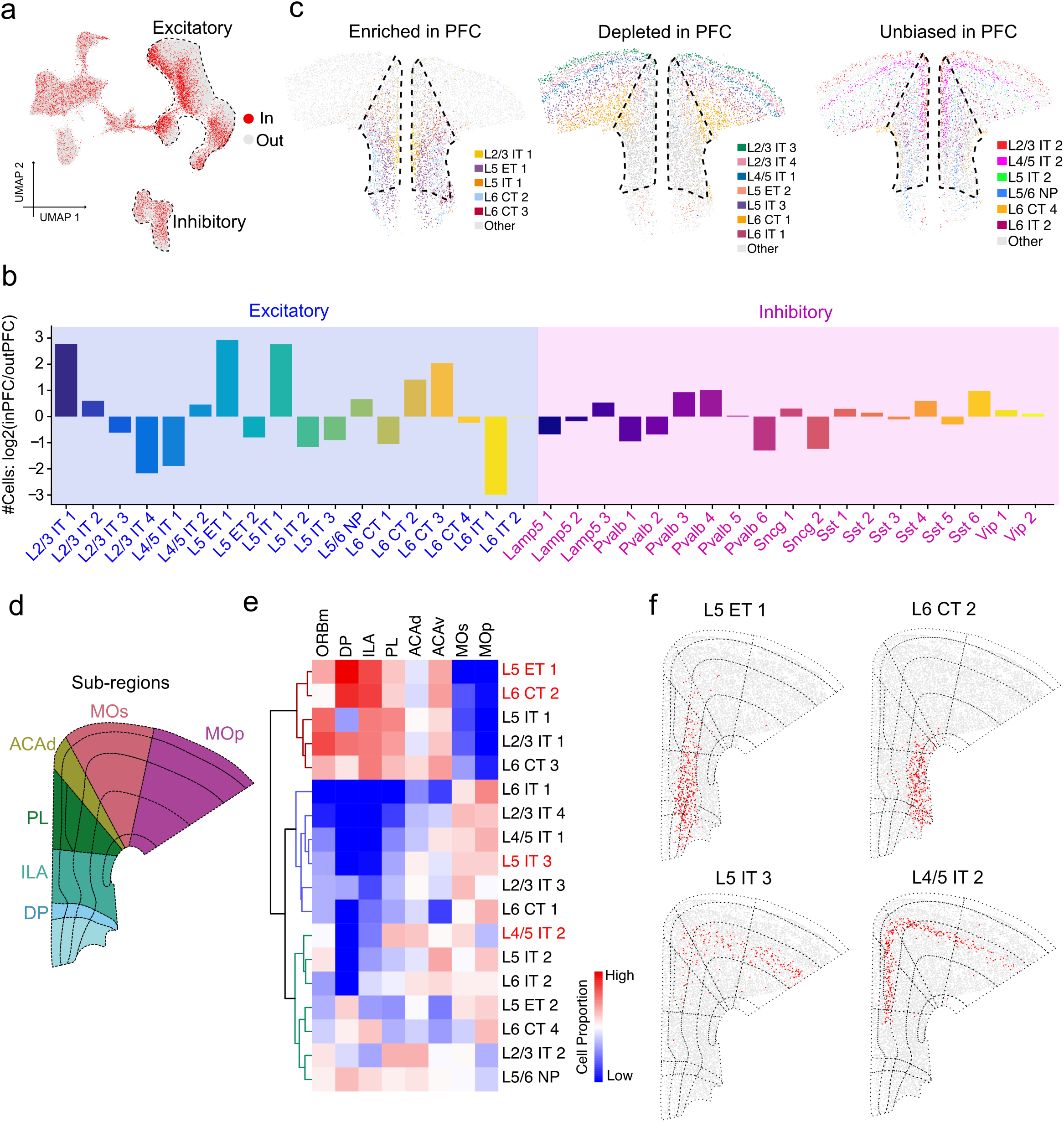
Distinct neuron subtypes are uniquely enriched or depleted in PFC relative to the adjacent cortical regions. **a**, UMAP of all MERFISH cells colored by their spatial location whether in or out of PFC. **b**, Barplot showing the log2 of the abundance ratio of subtype neurons in or out of PFC. **c**, Spatial location of excitatory neuron subtypes enriched (left), depleted (middle), and unbiased (right) distribution in PFC compare with adjacent regions. Dotted line marks PFC in the slice. **d,** Diagram of anatomical subregions of PFC and adjacent cortical regions. **e**, The normalized neuron proportion of excitatory subtypes in different anatomical subregions. **f**, Spatial location of four representative excitatory neuron subtypes on a coronal slice. Red dots represent the indicated subtypes. The dotted lines indicate the anatomical subregions from Allen Brain Atlas CCF v3.

The PFC has distinct functional subregions from its dorsal to ventral end, viz. anterior cingulate cortex (ACAd), prelimbic cortex (PL), infralimbic cortex (ILA) and dorso-peduncular cortex (DPP) (Fig. 3d). We asked whether these subregions have distinct cellular composition. Indeed, clustering with the normalized cell proportions across all subregions revealed the most enriched excitatory neurons in each subregion (Fig. 3e). Projecting cell types on to a coronal slice with subregion demarcations revealed heterogeneity of subtype distribution across different subregions (Fig. 3f). For example, L5 ET 1 is enriched in PL and ILA (but depleted in ACAd), while L6 CT 2 is mainly in ILA and L5 IT 3 is mainly in ACAd (Fig. 3e, f – enriched cells in 3e labeled red fonts). We also estimated the percent abundance of each cell type in each subregion (Fig. S5b). For example, L5 ET 1, L6 CT 2, L6 CT 3 or L6 CT 4 as well as inhibitory subtypes like Pvalb 5 or Sncg 2 are enriched in the infralimbic (ILA) relative to other subregions, while L2/3 IT 4, L4/5 IT 1, L5 IT 3 and especially L6 IT 1 are depleted in this subregion. Similarly, the dorsal anterior cingulate (ACAd) is enriched in L6 IT 1 or L2/3 IT 4, but depleted in L2/3 IT 1, L5 ET 2, L6 CT 2, L6 CT 3, L6 CT 4. Strikingly, the prelimbic (PL) maintains a steady share of cells from most subpopulations except for a higher percentage of L2/3 IT 2 and L4/5 IT 2. It is well established that many behavioral functions are specifically regulated by distinct subregions of the PFC. For example, conditioned fear response or trauma (as evidenced in PTSD) is encoded in the ILA^36^, while cue or context-associated reward memory is encoded in PL^37,38^, and compulsive behavior (often associated with drug addiction) is associated with ORBm^39^. Thus, revealing the differential neuron subtype distribution in the different PFC subregions may help link the PFC subregion-specific functions to the various differentially distributed neuron subtypes.

### Unique transcriptional signatures emerge in PFC

Functional differences across brain regions often underlie molecular adaptations^23^. The cortex is believed to be no exception. Thus, we asked whether the distinctive functions and cellular organization of the PFC is associated with specialized molecular features by comparing the transcriptome of PFC with that of the adjacent cortical regions. Indeed, a large number of genes interrogated in the MERFISH library are differentially expressed between the PFC and the neighboring cortices (Fig. 4a). Among the 416 genes analyzed, 54 were significantly enriched and 40 depleted in PFC (adjusted p-Value <0.05; DEG >20%) (Supplementary Table S2). Mapping expression of significantly enriched (*Nnat*) or depleted (*Scn4b*) genes onto the coronal section showed clear enrichment or depletion in the PFC region (Fig. 4b), which is consistent with the ISH data from the Allen Brain Institute (Fig. 4c), validating our MERFISH results.

**Fig. 4:**
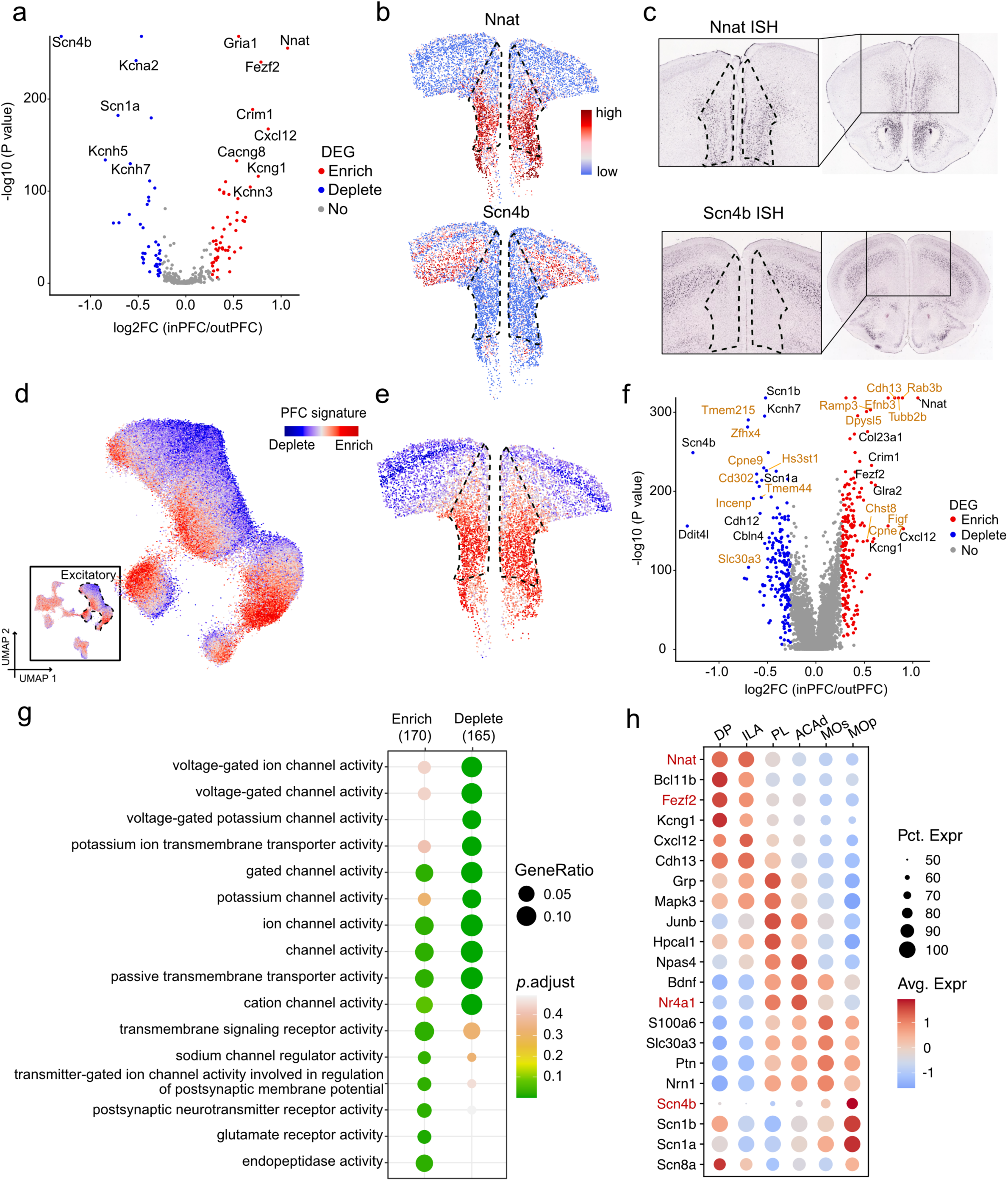
Genes with expression enriched or depleted in PFC. **a**, Volcano plot showing the differentially expressed genes (DEGs) that are enriched or depleted in PFC neurons relative to the neurons out of PFC. Expression of genes enriched, depleted in PFC are colored in red, blue dots, respectively. **b**, Spatial gene expression of *Nnat* (top) and *Scn4b* (bottom) in all excitatory neurons. Dotted line marks PFC region. **c**, *In situ* hybridization (ISH) data from Allen Brain Atlas showing the spatial expression of Nnat and Scn4b in a coronal slice (right) with zoom-in (left). **d**, UMAP of all MERFISH cells (bottom-left) and excitatory neurons colored by the PFC signature, which is defined as the average expression of top 10 enriched genes minus the average expression of top 10 depleted genes. **e**, Align the PFC signature onto a representative slice to show the spatial distribution of PFC signature. **f**, Volcano plot showing the expressions of genes enriched or depleted in PFC after imputing by iSpatial. A total 20,733 genes are analyzed. Genes analyzed by MERFISH are colored in black, and genes inferred by iSpatial are colored in yellow. **g**, The gene ontology enrichment analysis of genes that enriched or depleted in PFC. **h**, Gene expression enrichment analysis of genes enriched in the different anatomical subregions of PFC and the adjacent cortical regions.

We next asked whether specific types or categories of genes are selectively enriched or depleted in PFC. The differentially expressed genes had a strong representation of several ion channels and some key neurotransmitter receptors-which can impart very distinct electrical properties of the PFC relative to adjoining cortices^40^. In the ion channels group, several potassium channels are enriched or depleted (Supplementary table 2). The voltage-gated potassium channels subtypes^40–42^, especially delayed rectifiers (*Kcna2, Kcnb2, Kcnc2, Kcnc3, Kcnq3, Kcnq5*) are depleted (Fig. 4a, Supplementary table 2). However, some other delayed rectifiers (e.g., *Kcna1, Kcna4, Kcna5*, etc.) are not changed (Supplementary table S2). On the other hand, the inward (*Kcnh7)* and outward (*Kcnh5*) rectifier channels are depleted (Fig. 4a). Additionally, enrichment of BK channel like *Kcnmb4*, and reciprocally enriched modifier/silencer *Kcng1* (up), *Kcnf1* (up) but *Kcnv1* (down), were also observed (Fig. 4a, Supplementary table S2) Interestingly, *Kcnn3* is upregulated, that controls neuronal firing through after-hyperpolarization and its mutation is implicated in schizophrenia and bipolar disorder^43^. Apart from potassium, some prominent calcium channels (*Cacna1e, Cacna1h,* Fig. S6a) and sodium (*Scn3b*) channels, which have been implicated in major neurological disorders like autism and epilepsy^44–47^, are also enriched.

Apart from gated ion channels, another striking observation is the selective enrichment of *Gria1* (Fig. 4a), a principal ionotropic AMPA glutamate receptor subunit, in PFC. A GluA1 (protein product of *Gria1* gene) dimer binds a GluA2 (from *Gria2* gene) dimer to form a tetrameric ionotropic glutamate receptor. The GluA1 is strongly implicated in several neuropsychiatric disorders (schizophrenia, epilepsy, depression), chronic pain (increase) and drug addiction (decrease)^48^. Its expression in PFC declines with age and GluA1 is also implicated in Alzheimer’s disorder^49^. More interestingly, *Cacng8*, a transmembrane AMPA receptor-regulating auxiliary subunit, is also enriched within PFC (Fig. 4a). It regulates trafficking and gating of AMPA receptors and is implicated in several neuropsychiatric disorders (attention deficit or personality disorder)^50,51^. Enrichment of *Cxcl12* (Fig. 4a, S6a) is likely a final proof of the functional diversity of PFC. Aa a chemokine, *Cxcl12* plays roles from sculpting inhibitory neuron synapses to neuro-immune interactions, which in the adult cortex are characteristic of the PFC^52–54^.

To globally represent the remarkable transcriptional features of PFC neurons, we calculated the “PFC signature”, the average expression of the top 10 enriched genes minus top 10 depleted genes. When values for this index were projected (as red color) onto cells in the original UMAP, the PFC-enriched excitatory neurons clearly clustered and emerged (Fig. 4d). When PFC signature was mapped onto a representative coronal section, it localized precisely within the anatomical limits of the PFC (Fig. 4e), indicating a distinct molecular composition of the PFC relative to the adjacent cortices.

### iSpatial revealed transcriptome-wide PFC-enriched genes and functional pathways

To expand our spatial mapping of gene expression to the transcriptome scale, we next combined scRNA-seq with MERFISH to make predictions for spatial genes expression enrichment in PFC for genes not measured with MERFISH. To this end, we integrated our prior PFC scRNA-seq data^19^ and current MERFISH data to predict the expression pattern of all genes using iSpatial^55^, a bioinformatic tool we developed. The analysis revealed 190 PFC-enriched and 182 PFC-depleted genes (Fig. 4f, Supplementary table S2). Mapping enriched and depleted candidate genes predicted by iSpatial, *Cdh13* and *Abcd2* respectively, on to a coronal section revealed consistent localization with respect to the PFC boundaries (Fig. S6b), which is in line with the Allen Brain ISH results (Fig. S6b).

Gene Ontology enrichment analysis of the 364 spatially differentially expressed genes revealed *biological function* categories highly enriched in transporters, channels and receptor activity, which are known to modulate membrane potential (Fig. 4g). Depletion of voltage-gated potassium channels or transmembrane potassium transporter concur with a poised state of activity that PFC neurons must maintain for working memory function, a feature not essential for adjacent motor or sensory cortices^42,56^. Greater enrichment of ‘postsynaptic neurotransmitter activity’ or ‘glutamate receptor activity’ (Fig. 4g) relative to adjacent cortices reaffirm that PFC retains significant plasticity compared to these regions, even in adult. Curiously, some functions like ‘gated channel’ or ‘cation channel activity’ are both enriched and depleted (Fig. 4g). This indicates that PFC likely uses a different subset of receptors (class switching) for the same functions compared to adjacent cortices to adapt to its distinct electrophysiological needs.

A signaling pathways enrichment analysis of these 364 genes revealed opioid signaling, endocannabinoid pathway and glutamate receptor signaling as the top three pathways (Fig. S6c). While glutamate signaling is widespread in cortex, opioid and cannabinoid signaling are more uniquely characteristic of the PFC and are known to be essential for normal physiological functions of mood, memory, feeding, etc.^57–60^. This indicates that the distinct molecular composition of PFC is indeed tied to its specialized functions.

Decoding the transcriptome-wide, spatially enriched, gene expression patterns also allowed us to investigate whether there is expression bias between subregions of the PFC. Indeed, we detected several genes (e.g., *Nnat*, *Fezf2*, *Nr4a1*, and *Scn4b*, etc.) that are preferentially expressed in certain subregions of the PFC (Fig. 4h, S6d), which are also validated in Allen Brain ISH data (Fig. S6d). Thus, subregion-specific functions of PFC are potentially enabled by their discrete molecular compositions imparting specific electrical and signaling properties.

### Spatial organization predicts subtype-specific interactomes in PFC

Extensive local processing of convergent and divergent signals is one of the principal characteristics of cortex and is particularly prominent in PFC, an area that integrates sensory/cognitive inputs in real time to govern executive function^61^. Integration of multilevel (thalamic, cortical or subcortical) inputs, their transfer through cortical layers, and modulation by interneuron inhibition as well as disinhibition all rely on extensive local interactions between neurons that are neighboring or located in close proximity within PFC^34,62–64^. Given that MERFISH allowed the mapping of the precise location of every cell in PFC, we explored the potential cell-cell interactions at the cell subtype level. To this end, we inspected the cell subtypes composition of the neighboring cells for each cell and calculated the enrichments of paired subtype-subtype colocalizations. Enrichment of proximity was notable amongst many groups of cells (Fig. S7a). IT subtypes of L2/3 are closely apposed in the superficial layers and engage in cortico-cortical interactions to integrate signals from sensory and association cortices (Fig. S7a). Interestingly, most of these subtypes have interactions with L4/5 IT subtypes (Fig. S7a) that receive exclusive inputs from thalamus or lower order cortex (since PFC has no clear L4), and are known to relay processed information mainly to L2/3^65^. This observation reinforces the notion that spatial organization of neuronal types reflect the order of information flow within the circuits they comprise, which in turn emerges as the systematic layered cortical structure. Interestingly, our analysis revealed specific interactions in the deeper layers that may not be apparent from histological organization alone. For example, L6 IT neurons (like L6 IT 1) share proximity with specific ET neurons (L5 ET 2), revealing subtype selectivity (and in turn circuit selectivity) within L5-L6 communication (Fig. S7a). Subtype selectivity is perhaps most important in excitatory-inhibitory coupling. The inhibitory *Pvalb* neurons directly access the soma of excitatory pyramidal neurons to regulate firing through feed forward inhibition^66^. Preferential pairing of many excitatory subtypes with one or few (but not all) specific Pvalb subtypes were detected in our analysis (Fig. S7a). For example, L5 IT 3 scored the highest proximity with Pvalb 1, while L5 ET 2 (located within the same layer) has greater interaction probability with Pvalb 6 (Fig. S6a-highlighted boxes). Mapping cells on to a representative coronal section revealed the relative proximities of each of these two excitatory-inhibitory pairs, and also a different spatial enrichment of the Pvalb 1 and Pvalb 6 subtypes (Fig. S7b).

In summary, MERFISH measurements in the PFC allowed us to provide an entry point (or repository) for predicting subtype-specific synaptic interactions in the 3D anatomical space, which can be then studied by using appropriate experimental approaches.

### Spatial and molecular organization of PFC projection circuits to major subcortical targets

It is well known that the PFC excitatory pyramidal neurons project to different subcortical targets including striatum, nucleus accumbens, thalamus, hypothalamus, amygdala, periaqueductal gray or ventral tegmental area^18,67^. However, the spatial organization and whether different neuron subtypes project to different targets are not well characterized.

A prior study has performed retrograde labeling and scRNA-seq for some of these major targets of labeled PFC neurons^2^. We integrated our PFC MERFISH data with this dataset to predict the PFC neuron subtypes with spatial/layer location projecting to these different targets. Through joint embedding and supervised machine learning, we could assign respective projection identity to the molecular clusters organized in space within the PFC (Fig. 5a). An overlap of the MERFISH and scRNA-seq clusters through UMAP visualization revealed a strong correspondence (Fig. 5b, S8a). The ROC curve for the prediction model independently predicted 6 different projection targets with high confidence, including contralateral PFC (cPFC), dorsal striatum (DS), hypothalamus (Hypo), nucleus accumbens (NAc), periaqueductal gray (PAG), and amygdala (Amyg) (Fig. 5c). Mapping these projection cells onto a coronal slice of frontal cortex revealed the identity and spatial organization of neurons that project to each of these 6 targets within the PFC (Fig. 5d). Distinct spatial localization of each of these 6 groups of cells can be visualized when mapped individually on the coronal slice (Fig. S8b). This analysis allowed us to associate different subsets of each neuronal type that project to different regions with their location within the brain (Fig. 5e), which reveals that most of the target brain regions receive projection from more than one neuron subtypes. For example, the amygdala receives projections from all four subtypes of L6 CT neurons as well L5 ET 1 neurons, but the majority comes from L6 CT 2. Likewise, the hypothalamus receives its projections from L5 ET 1 and L6 CT 1; dorsal striatum from L6 CT 1, 2, and 3; and NAc gets mainly from L6 CT 1, L5 ET 1 and some from L6 CT 2. However, one exception to the general rule is the PAG, which receives its projections almost exclusively from L5 ET, predominantly from L5 ET 1 (and some from L5 ET 2). Consistent with prior knowledge, superficial layer IT neurons project to the contralateral hemisphere of PFC^68^. It should be noted that each of these target brain regions are involved in many different behavioral functions. For example, amygdala alone is implicated in fear, addiction, emotion, memory and pain^69–71^. It is expected that inputs will likely to be received from diverse projections (of different neuron subtypes) to selectively trigger specific synapses/pathways to sustain the functional complexity of PFC.

**Fig. 5:**
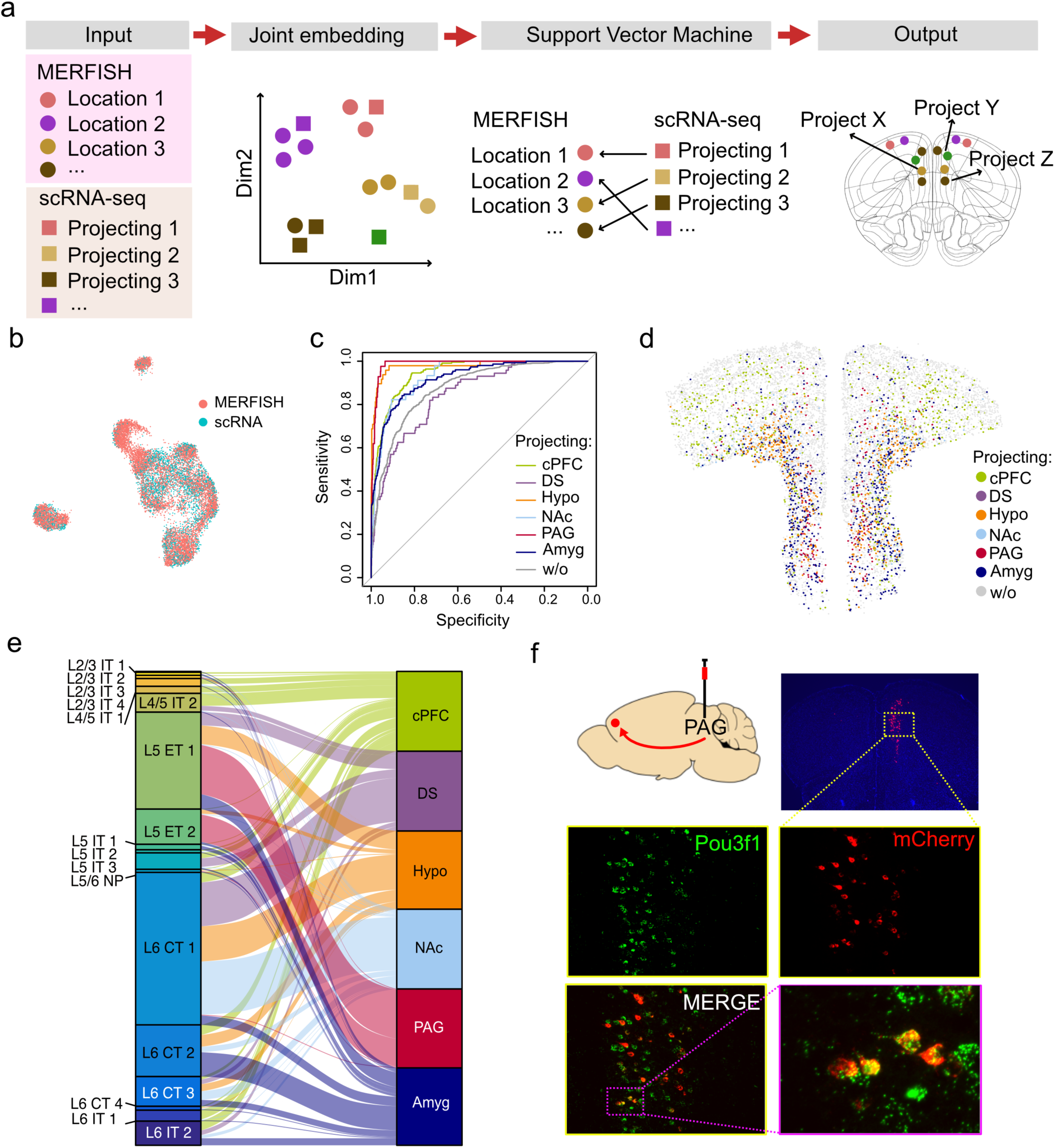
Spatial and molecular organization of PFC excitatory subtypes projection to the major PFC targets. **a**, Schematics of the strategy for inferring neuronal projection of MERFISH clusters. The MERFISH and scRNA-seq data are integrated into a reduced dimensional space. A support vector machine is used to predict neuronal projection of the MERFISH neuron subtypes (see methods). **b**, UMAP visualization of cells derived from MERFISH and scRNA-seq data after integration. **c**, The ROC curves showing the prediction powers of six projection targets. w/o represents the cells without projection information. **d**, A coronal slice showing *in silico* retrograde tracing from six injection sites, labeled by different colors as indicated. **e**, The inferred projection targets of molecularly defined excitatory neuron subtypes, represented by an alluvial diagram. **f**, PFC to PAG projection validation. Retrograde mCherry expressing AAV was injected in PAG and brain slice of PFC was used for smFISH. mCherry (red) labeled neurons co-express the L5 ET1 marker *Pou3f1* (green). All mCherry positive neurons are Pou3f1-positive.

To validate our computational model-based neuron projection predication, we performed retrograde tracing from two of these target regions by injecting retrograde AAV virus into the PAG and the amygdala to drive mCherry expression. Four weeks after the injection, we prepared cryosections and performed single molecule FISH (RNAScope) to co-label mCherry RNA and the respective cell-type specific markers. Consistent with the prediction, all mCherry expressing PAG retro-traced PFC neurons exclusively colocalized with *Pou3f1,* a selective marker for L5 ET 1 and L5 ET 2 (Fig. 5f). In contrast, colocalization of mCherry was detected for both *Pou3f1* (L5 ET 1) and *Foxp2* (L6 CT) for amygdala as predicted (Fig. S8c). These data support the accuracy of our circuit predications.

### Identifying PFC neuron subtypes involved in chronic pain

Functions of PFC in cognition or execution are most widely studied. However, besides those voluntary behaviors, PFC also plays a pivotal role in autonomically modulating pain perception, and aberrations in this process is emerging as a major player in pain “chronification”^5,14^. While chronic pain is escalating as a leading healthcare challenge^7^, molecular underpinnings of the dysfunction remain unknown. Chronic pain has been strongly associated with transcriptional adaptations across the PFC^5,72,73^, the spatial or cell type-specific resolution of these changes are less clear. To explore the utility of our MERFISH datasets, we attempted to identify the PFC neuron subtypes involved in chronic pain by identifying the neuron subtypes that undergo the strongest transcriptional response in chronic pain.

To this end, we utilized the well-established spared nerve injury (SNI) model of chronic neuropathic pain in mice^74^ where two of the three branches of the sciatic nerve are transected (Fig. 6a), which causes a state of chronic neuropathic pain in hind paw that lasts for months. Six weeks after surgery, brains from 3 pairs of sham and SNI mice were collected and characterized with MERFISH (Fig. 6a). Of all the neuronal subtypes, L5 ET 1 registered the strongest transcriptional response (with largest total number of differentially expressed genes), followed by the L6 CT 2 and L5 ET 2 (Fig. 6b). Lesser changes were detected in L6 CT 3, L5 IT 1, L6 IT 2, L2/3 IT 2 and L4/5 IT 2 (Fig. 6b). No significant changes were detected in the other 30 clusters despite many of the excitatory neuron subtypes being highly abundant in PFC, suggesting these clusters are minimally affected in chronic pain. Interestingly, the two highest impacted clusters respectively project to PAG (L5 ET 1) and amygdala (L6 CT 2) (Fig. 5e), the two major hotspots known to regulate sensory and affective aspects of pain^5,71^.

**Fig. 6:**
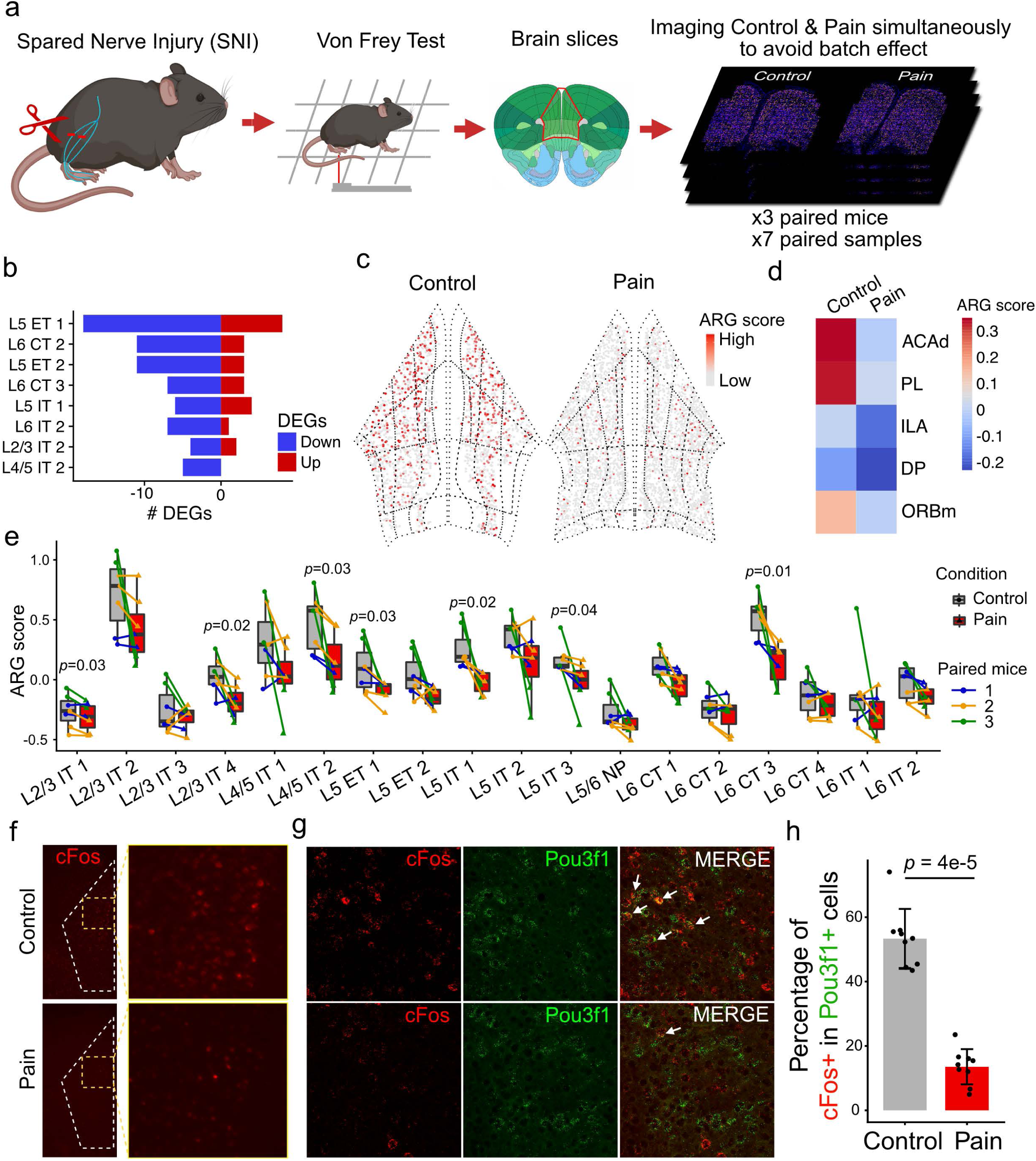
Chronic pain caused cellular and molecular changes in PFC excitatory neurons. **a**, Overview of chronic pain sample preparation. For each MERFISH run, one brain slice from each of control and pain condition are loaded together to avoid batch effect. Seven paired samples from three paired mice were imaged. **b**, The numbers of differentially expressed genes comparing pain and control samples for the indicated neuron subtypes are shown. The numbers of up-regrated and down-regulated genes are colored in red and blue, respectively. **c**, Spatial distribution of cells colored by activity-regulated genes (ARG) scores in control and pain conditions. The anatomical subregions of PFC are also shown. **d**, Heatmap showing ARG score in PFC subregions in pain and control samples. **e**, ARG scores of PFC excitatory subtypes in pain and control samples. Paired dots represent the control-pain paired samples which were imaged together. Color of the paired dots represent the paired mice ID. Two-tailed paired t-test is used to calculate the p-value. **f,** Global overview of PFC in half coronal section with Fos smFISH (red) in Sham (Control) and chronic pain conditions. **g,** smFISH co-labeling of Fos and Pou3f1 (L5 ET marker) at high magnification in Sham and chronic pain conditions. Arrowheads in merged images indicate double positive neurons. **h**, Barplot showing the percentage of cFos+ cells to Pou3f1+ cells. Nine random fields are surveyed. Two-tailed Mann-Whitney test is used to calculate the p-value.

Chronic pain is known to inflict strong and sustained hypoactivity across the PFC^9,12,14,75^. We asked whether this can be detected in the baseline expression of neuronal activity-regulated genes (ARGs) to identify prominently affected neuron subtypes. We calculated ARG score using the mean expression of a panel of 5 ARGs (Arc, Junb, Fos, Npas4, Nr4a1) and compared between sham and SNI groups. We observed a strong and widespread reduction of ARG score when it is plotted on representative coronal sections (Fig. 6c). A subregion-specific calculation revealed that the ACAd and PL are the most impact PFC regions (Fig. 6d). We next compared the differences of ARG score across the individual excitatory neuron clusters (Fig. 6e), and found it is downregulated in several clusters, including those exhibiting transcriptional changes (e.g., L5 ET 1, L6 CT 3, Fig. 6b).

To validate chronic pain-induced hypoactivity across PFC, we performed single molecule FISH (smFISH) to compare Fos expression between sham and SNI brain sections. Although sham shows a baseline Fos activity in PFC, a general Fos depletion is obvious in the SNI (Fig. 6f). Co-staining Fos with Pou3f1, a selective marker for L5 ET1, revealed significant Fos depletion in this neuron subtype in the SNI brains (Fig. 6g, h).

Despite the conventional knowledge that a PFC-PAG circuit is involved in descending modulation of pain^5^, its cell type identity or changes in chronic pain were unclear. Our findings revealed the molecular identity and spatial organization of this circuit: the L5 ET 1 neurons with PAG projection (Fig. 5e), which are strongly deactivated in chronic pain (Fig. 6g) with the maximum transcriptional adaptation (Fig. 6b). Additionally, we also identified at least two CT subtypes in L6 (L6 CT 2 and 3) that project to limbic structures like amygdala, NAc and hypothalamus (Fig. 5e) that may be involved in the affective response to pain.

## Discussion

In this study we present an account of how the PFC is distinctly organized at the cellular, molecular and projection levels relative to the adjacent regions within the frontal cortex. We exploit this characterization to reveal the molecular identity of key neuron subtypes that are engaged in chronic pain, and, more broadly, we provide a resource for the systematic mapping of functional ensembles and circuits selectively engaged in various cognitive and executive functions associated with PFC. Spatial transcriptomics is a rapidly growing field^76^ and similar to recently reported brain regions^29,32,77^, MERFISH enabled a systematic decoding of PFC’s cellular and molecular organization.

### The diversity of PFC neuron subtypes is consistent with its functional diversity

We observed that there were a variety of neuronal subtypes largely specific to the PFC or to surrounding regions (Fig. 3a-c). Cellular composition of a cortical area should be predominantly governed by the input and output circuits associated with its function. This regional neuronal subtype specificity, in turn, may underlie the unique properties of the PCF relative to other cortical regions. For example, the PCF is agranular and lacks a typical L4, associated with thalamic input, it receives long-range inputs across all of its layers, and PFC neurons project to subcortical targets from almost all layers while PFC neurons engage in reciprocal connections with most of these functions ^78,79^. Perhaps for the diverse functions, there is a 2-fold enrichment of the superficial-most IT neurons (L2/3 IT 1) to handle the cortico-cortical communications, but the subsequent IT populations (L2/3 IT 4 or L4/5 IT 1) are markedly depleted to make room for enrichment of L5 IT 1 or L5 ET 1 that engage in long distance subcortical projections. Enrichment of two CT subtypes (L6 CT 2 and 3) is consistent with the observation that CT neurons of PFC, unlike other cortices, project to several subcortical targets (Fig. 5e), rather than thalamus alone. Notably, two of these enriched neuron subtypes (L5 ET 1 and L6 CT 2) eventually emerge as key subtypes engaged in chronic pain, a function exclusively performed by PFC (Fig. 6).

Depletion of certain subtypes of Pvalb (Pvalb 1, 2, 6), which also accounted for an overall lower count of Pvalb neurons in the PFC (relative to the adjacent regions), suggests that feedforward inhibition is differently organized in PFC. This either indicates an overall lower level of feed forward inhibition and perhaps a greater flexibility in excitatory-inhibitory balance; or larger receptive fields are covered by individual Pvalb neurons, synapsing with more pyramidal neurons towards a goal of regional synchronization. In either case, this is an important observation as functional imbalance of Pvalb neurons has been implicated in almost every PFC-associated diseases, such as schizophrenia^80^, bipolar, depression, and chronic pain^81^. It should be noted that detection of such regional differences would not be possible without the spatial profiling techniques like MERFISH.

Besides cellular composition, we detected strong transcriptional features unique to the PFC. We found expression of a large number of ion channels and receptors is selectively increased or decreased in PFC relative to the adjacent cortical regions (Fig. 4). It is generally appreciated that different cortical regions have different baseline electrical properties and qualitatively different activity patterns, which in turn is critical for its specific function^23^. For example, sensory cortices, such as visual cortex, have millisecond scale dynamics which is believed to be much faster than that of frontal regions involved in decision, deliberation or short-term memory. Recording of electrical field potentials across cortical areas provide strong evidence supporting such regionally variable activity patterns^82,83^. However, the biological substrates underlying such functional differences have been less clear. Our findings revealing preferential expression or repression, or even subtype switch of a wide range of cation channels and key glutamate receptor subunits in PFC establish a foundation for identifying the potential biological substrates explaining the diverse PFC functions.

### The diverse projections of the PFC neuron subtypes

As the apex controlling center for cognitive, executive and emotional behaviors, PFC has one of the most diverse efferent projection profiles amongst all cortical areas. However, a striking observation was that while targets like PAG receives projection from a more homogeneous molecular subtype in L5 (L5 ET), most other brain regions receive heterogeneous projections from multiple cell types of different layers (Fig. 5). Although intriguing, this may in fact reflect a more sophisticated model of top-down control by PFC. Most of these target regions such as amygdala, striatum, nucleus accumbens, and hypothalamus engage in many different behavioral processes, which may also be regulated by distinct groups of neurons within each of these target regions. Accordingly, different lines of afferent projections from PFC can synapse on to different neuron populations to form separate circuits within a target and thereby separately modulate different behaviors under different contexts. Additionally, different subtypes in separate layers can receive distinct upstream inputs within PFC that can be separately relayed to the targets through specific projection clusters. For example, L5 ET1 and L6 CT2 may receive different upstream inputs in PFC and can also project to separate cell types within amygdala to modulate different behaviors in separate contexts (Fig. 5e). Further work in this direction should reveal the cellular and molecular organization of all PFC projection circuits and identify specific ensembles engaged by different behaviors.

### The L5 ET1 neuron subtype might regulate chronic pain through PFC to PAG projection

We identified the key cell types that are specifically impacted in PFC under chronic pain (Fig. 6). Amidst the rising prevalence of chronic pain and emerging consensus that transition to chronic pain is centrally regulated, there has been little clarity about the cellular and molecular mechanisms underlying the chronification, which is key to therapeutic targeting. Previous studies have shown that transcranial stimulation of PFC could relieve chronic pain^15,84,85^. Such studies, although established a causal connection, did not provide a long-term solution for pain management owing to the deleterious effects of broad non-specific cortex-wide stimulations. Despite a long-standing knowledge of putative PFC to PAG projections in descending inhibition of pain^5^, the molecular identity of this circuit was unknown. In this regard, our study revealing the L5 ET 1 as a major neuron subtype with exclusive projection to the PAG, and undergoes transcriptional changes under chronic pain state is of particular relevance. While it is likely that deactivation of this cluster will impair descending inhibition of pain which paves way for persistent pain/sensitivity, it remains to be determined if it also contributes to the affective component of pain. However, L6 CT 2 and L6 CT 3, the two other implicated clusters, project to multiple limbic regions including amygdala, NAc and hypothalamus, and their dysfunctions may elicit strong negative effect characteristic of chronic pain states^71,86,87^. All these remain valuable prospects for future functional studies through targeted neuronal activity manipulation using genetically engineered animal models.

## Materials and methods

### Mice and Surgery

All experiments were conducted in accordance with the National Institute of Health Guide for Care and Use of Laboratory Animals and approved by the Institutional Animal Care and Use Committee (IACUC) of Boston Children’s Hospital and Harvard Medical School. Wildtype male C57BL6 mice of about 10 weeks old were used for the study. Mice were maintained at 12h light/dark cycles with food and water ad libitum. For the spared nerve injury surgery, mice were anesthetized with ketamine. Hair was shaved above the knee on one side (usually left) and the skin was sterilized with iodine and isopropanol. The muscles were separated by blunt dissection to expose all three branches of the sciatic nerve. The tibial and common peroneal branches of the nerve that run parallel were tied tightly with two sutures and a piece between the two ties was transected and removed. Care was taken that the third branch (sural nerve) was untouched during the whole procedure. The retracted muscles were released, and the skin stitched back. In the sham surgery group, identical steps were followed to expose the nerve, but no transection was performed, and skin was stitched back in position. Mice tissues were harvested 6 weeks after the surgery.

### MERFISH Encoding Probes

A library of MERFISH encoding probes for all target genes was generated as described previously^29^. Briefly, a unique binary barcode was assigned to each gene based on an encoding scheme with 24-bits, a minimum Hamming distance of 4 between all barcodes, and a constant Hamming weight of 4. This barcoding scheme left 60 ‘blank’ barcodes unused to serve as a measure of false-positive rates. For each gene, 50 to 70 30-nt-long targeting regions with limited homology to other genes and narrow melting temperature and GC ranges were selected, and individual encoding probes to that gene were created by concatenating two 20-nt-long readout sequences to each of these target regions. Each of the 24 bits were associated with a unique readout sequences, and encoding probes for a given gene contained only readout sequences for which the associated bit in the barcode assigned to that gene contained a ‘1’. Template molecules to allow the production of these encoding probes were designed by adding flanking PCR primers, with one primer representing the T7 promoter. This template oligopool was synthesized by Twist Biosciences and enzymatically amplified to produce encoding probes using published protocols^29^.

### MERFISH tissue processing and imaging

Tissue was prepared for MERFISH as described previously^29^. Briefly, mice were euthanized under CO_2_ and brains were quickly harvested and rinsed with ice-cold calcium and magnesium free PBS. The brains were frozen on dry ice and stored at −80 ^0^C till sectioning. The frozen brains were embedded in OCT on a mixture of ethanol and dry ice. Serial 14-μm-thick sections of the frontal cortex spaced about 150 μm apart were collected and placed on poly-D lysine coated, silanized coverslips, containing orange fiducial beads, prepared as described previously^29^. The sections were allowed to briefly air dry and immediately fixed with 4% PFA for 10 mins. Sections were washed in PBS and stored in 70% ethanol for at least 12h to permeabilize. The sections were washed in hybridization buffer (2xSSC+30% formamide) and then drained and inverted over parafilm in petri dish onto a 50 μl droplet of mixture containing encoding probes and a poly(A) anchor probe^29^ in hybridization buffer (2xSSC, 30% formamide, 0.1% yeast tRNA, 10% dextran sulfate) and hybridized in a covered humid incubator at 37^0^C for 2 days. Coverslips were then washed in hybridization buffer and the sections were embedded into a thin film of poly-acrylamide gel, as described previously. The embedded sections were then digested for 2 days in a 2xSSC buffer containing 2% SDS, 0.5% Triton X-100 and 1:100 proteinase K. The coverslips were washed and stored in 2xSSC at 4 ^0^C until imaging. MERFISH imaging was performed on a custom microscope and flow system, as described previously^29^. In each imaging round, the volume of each slice was imaged by collecting a z-stack at each field-of-view containing 10 images each spaced by 1 micron. 12 imaging rounds using two readout probes per imaging round were used to read out the 24-bit barcodes. Readout probes were synthesized by Biosynthesis and contained either a Cy5 or Alexa750 conjugated to the oligonucleotide probe via a disulfide bond, which allowed reductive cleavage to remove fluorophores after imaging, as described previously. A readout conjugated to Alexa488 and complementary to a readout sequence contained on the polyA anchor probe was hybridized with readouts associated with the first two bits in the first round of imaging.

### Image processing, decoding and cell segmentation

MERFISH data was decoded as previously described^29^. Briefly, images of fiducial beads collected for each field-of-view in each imaging round were used to align images across imaging rounds. RNAs were detected using a pixel-based approach, in which images were first high-pass filtered, deconvolved, and low-pass filtered. Differences in the brightness of different imaging rounds were corrected by an optimized set of scaling values, determined from an iterative process of decoding performed on a randomly selected subset of fields-of-view, and the intensity trace for individual pixels across all imaging rounds was matched to the barcode with the closest predicted trace as judged via a Euclidean metric and subject to a minimum distance. Adjacent pixels matched to the same barcode were aggregated to form putative RNAs. RNA molecules were then filtered based on the number of pixels associated with each molecule (greater than 1) and their brightness to remove background.

As described previously^29^, the identification of cell boundaries within each FOV was performed by a seeded watershed approaching using DAPI images as the seeds, and the poly(A) signals to identify segmentation boundaries. Following segmentation, individual RNA molecules were assigned to specific cells based on localization within the segmented boundaries.

### Preprocessing of MERFISH data

The decoded data was preprocessed by the following steps: 1) Segmented “cells” with a cell body volume less than 100 µm^3^ or larger than 4000 were removed; 2) Cells with total RNA counts of less than 10 or higher than 98% quantile, and cells with total RNA features less than 10, were removed; 3) To correct for the minor batch fluctuations in different MERFISH experiments, we normalized the total RNA counts per cell to a same value (500 in this case); 4) Doublets were removed by Scrublet^88^; 5) The processed cell-by-gene matrix was transferred to gene-by-cell matrix and then loaded into Seurat V4^89^ for downstream analysis. The matrix was log-transformed by the Seurat standard pipeline.

### Cell clustering

Two rounds of cell clustering were used to identify cell types and subtypes. In the first round, we identified the three major cell types: excitatory neurons, inhibitory neurons, and non-neuronal cells. In the second round, each major cell type was further clustered. Excitatory neuron was further clustered into 18 subtypes, inhibitory neurons was further clustered into 19 subtypes, non-neuronal cell was further clustered into 15 subtypes. Then, we separated the excitatory subtypes into seven groups according to the neuronal projection: L2/3 IT, L4/5 IT, L5 IT, L6 IT, L5 ET, L5/6 NP, and L6 CT. The inhibitory neuron was cataloged into five groups based on the main markers: Lamp5, Pvalb, Sncg, Sst, and Vip. Non-neuronal cells were cataloged into six groups: endothelial cells, microglia, oligodendrocytes, OPC, astrocytes, and VLMC. Each round of clustering following the same workflow as described previously. First, all gene expression was centered and scaled via a z-score, and PCA was applied on the scaled matrix. To determine the number of principal components (PCs) to keep, we used the same method described before^29,77^. Briefly, the scaled matrix was randomly shuffled and PCA was performed based on the shuffled matrix. This shuffling step was repeated 10 times and the mean eigenvalue of the first principal component crossing the 10 iterations was calculated. Only the principal components derived from the original matrix that had an eigenvalue greater than the mean eigenvalue were kept. Harmony^90^ was then used to remove apparent batch effect among different MERFISH samples. The corrected PCs were used for cell clustering. The nearest neighbors for each cell were then computed by a K-nearest neighbor (KNN) graph in corrected PC space. Bootstrapping was used for determining the optimal k value for KNN as previously described^29,77^ (k = 10 in the first round clustering. k = 50, 20, 15 for excitatory neurons, inhibitory neurons, and non-neuronal cells in the second round). Leiden method was used for detecting clusters^91^. The resolution was set to 0.3 in the first round clustering, and to 2 for the second round. Finally, we manually removed the clusters representing doublets, which express high levels of the established markers of multiple cell types. Clusters located outside of the cortex were also removed.

### Correspondence between scRNA-seq and MERFISH clusters

To compare the cell clusters identified by scRNA-seq and MERFISH, we first co-embedded the two datasets in a corrected PCA space using Harmony as described above. Then, all the cells from both scRNA-seq and MERFISH were used to build the KNN graph. The first 30 corrected PCs were inputted into Seurat::FindNeighbors to compute the KNN. For each cell cluster in MERFISH, we obtained the cell cluster’s nearest 30 neighbor cells’ information. Then, we calculated the percentages of the cell clusters derived from scRNA-seq that were near to this MERFISH cluster, from which we obtained a correspondence matrix, where each row is a cluster from scRNA-seq, each column is a cluster from MERFISH, the element in the matrix indicates the similarity between the two clusters. Similarly, for each cell cluster in scRNA-seq, we inquired the nearest clusters derived from MERFISH data to generate another correspondent matrix. The average of the two correspondent matrices were used to indicate the similarities between the cell clusters defined by scRNA-seq and MERFISH.

### Cell-cell proximity

For each cell, we first identified the nearest 30 neighbors based on spatial distance. Next, we derived the cell subtypes of these neighboring cells, and obtained the cell subtypes composition of these cells nearby the inquired cell. After iteration of all cells in all subtypes, we could calculate the number of occurrences of paired cell-cell and obtain the cell-cell proximity matrix (Observed matrix). Because of the cell number differences for each subtype, we normalized cell-cell proximity matrix by a random shuffled matrix (Expected matrix). To derive the shuffled matrix, we first shuffled the cell identities by random assign a subtype for each cell. Then, the random cell-cell proximity matrix was calculated by the same method before. Finally, the normalized cell-cell proximity matrix was calculated by log2(Observed matrix/Expected matrix). In addition, the p-values were calculated by wilcoxon rank tests (using wilcox.test in R) and then adjusted by Benjamini-Hochberg method (using p.adjust in R, method = “BH”).

### Excitatory neuron projection prediction

The scRNA-seq data (GEO: GSE161936)^2^ was first preprocessed by standard Seurat pipeline. Only the cells from dorsomedial (dmPFC) and ventromedial (vmPFC) regions were used. We integrated the MERFISH and scRNA-seq data using Harmony, and all the cells derived from MERFISH/scRNA-seq were co-embedded on a corrected PCA space. The first corrected 30 PCs were selected as features to train a multi-class support vector machine (SVM) for predicting the neuronal projection. The cells from scRNA-seq were separated into training and test groups. Then, the SVM was trained on training data and validated on test data by using the radial basis function kernel. Gamma was set to 0.01, cost was set to 10. The receiver operating characteristic (ROC) curve was plotted to evaluate the performance using pROC package in R and the area under the AUC curve (AUC) was equal to 0.913. Finally, the model was applied to MERFISH cells to predict their projections.

### Register MERFISH slice to Allen Brain Atlas

To align MERFISH slices to the Allen common coordinate framework (CCF) v3 we leveraged the spatial distribution of cells identified by MERFISH in each slice as well as DAPI images of that slice. First, each brain slice was paired to the closest matching coronal section in CCF v3 with the help of DAPI image and spatial location of the cell types. Then, we modified the WholeBrain package^92^ to align the MERFISH slice to the corresponding matching CCF coronal section. To assure accurate alignment, we leveraged the MERFISH cell typing result at single cell resolution and used certain cell types as anchors to help locating the anatomic features. VLMC cells are used for marking the surface of brain slice as follows: Inhibitory neuron subtype, Lamp5 3, for locating layer 1; L2/3 IT neurons for locating layer 2, L6 CT neurons for locating layer 6, oligodendrocytes for locating corpus callosum. Since some small slices do not have sufficient features to align, 45 out of 60 slices are successfully registered to CCF v3, which allowed us to define the anatomic PFC and PFC subregions.

### Differentially expressed genes between pain and control conditions

To detect differentially expressed genes (DEGs) and correct the batch effects, we used a logistic regression framework. For each gene, we constructed a logistic regression model to predict the sample conditions *C* by considering the batch information *S*, *C* ~ *E* + *S*, and compared with a null model, *C* ~ 1 + *S*, with a likelihood ratio test. Then, Bonferroni correction method was applied to adjust for multiple comparisons. Here, “LR” method in Seurat FindAllMarkers was used for conducting this analysis.

### Data and code availability

The MERFISH data generated in this study has been deposited to Brain Image Library with accession number: in the process of uploading. Interactive visualization of MERFISH data can be accessed at: https://yizhang-lab.github.io/PFC. Code for MERFISH analysis is available at https://github.com/YiZhang-lab/PFC-MERFISH.

## Acknowledgements

We thank Drs. Barbara Calderone and Olga Alekseenko from the Harvard Neurobehavior Core for their support; Dr. Renchao Chen for critical reading of the manuscript and valuable suggestions; Drs. Yingying Zhang, Yuanyou Wang and Paolo Cadinu, and Ms. Rosalind Xu for help with MERFISH troubleshooting and pipelines. This project was supported by the National Institutes of Health (1R01DA042283, 1R01DA050589), the Open Philanthropy Foundation, and the HHMI. Y.Z. is an investigator of the Howard Hughes Medical Institute. This article is subject to HHMI’s Open Access to Publications policy. HHMI lab heads have previously granted a nonexclusive CC BY 4.0 license to the public and a sublicensable license to HHMI in their research articles. Pursuant to those licenses, the author-accepted manuscript of this article can be made freely available under a CC BY 4.0 license immediately upon publication.

## Author Contributions

Y.Z. conceived the project; A.B., Y.Z., and J.R.M. designed experiments; Y.Z. and J.R.M. supervised the project; A.B. performed experiments; C.Z. analyzed the data; M.N.D helped with probe design; B.W. built the MERFISH microscope and trained A.B. for MERFISH experiments; A.B., C.Z., Y.Z. and J.R.M. interpreted the data and wrote the manuscript.

## Supplementary figure legends

**Fig. S1:**
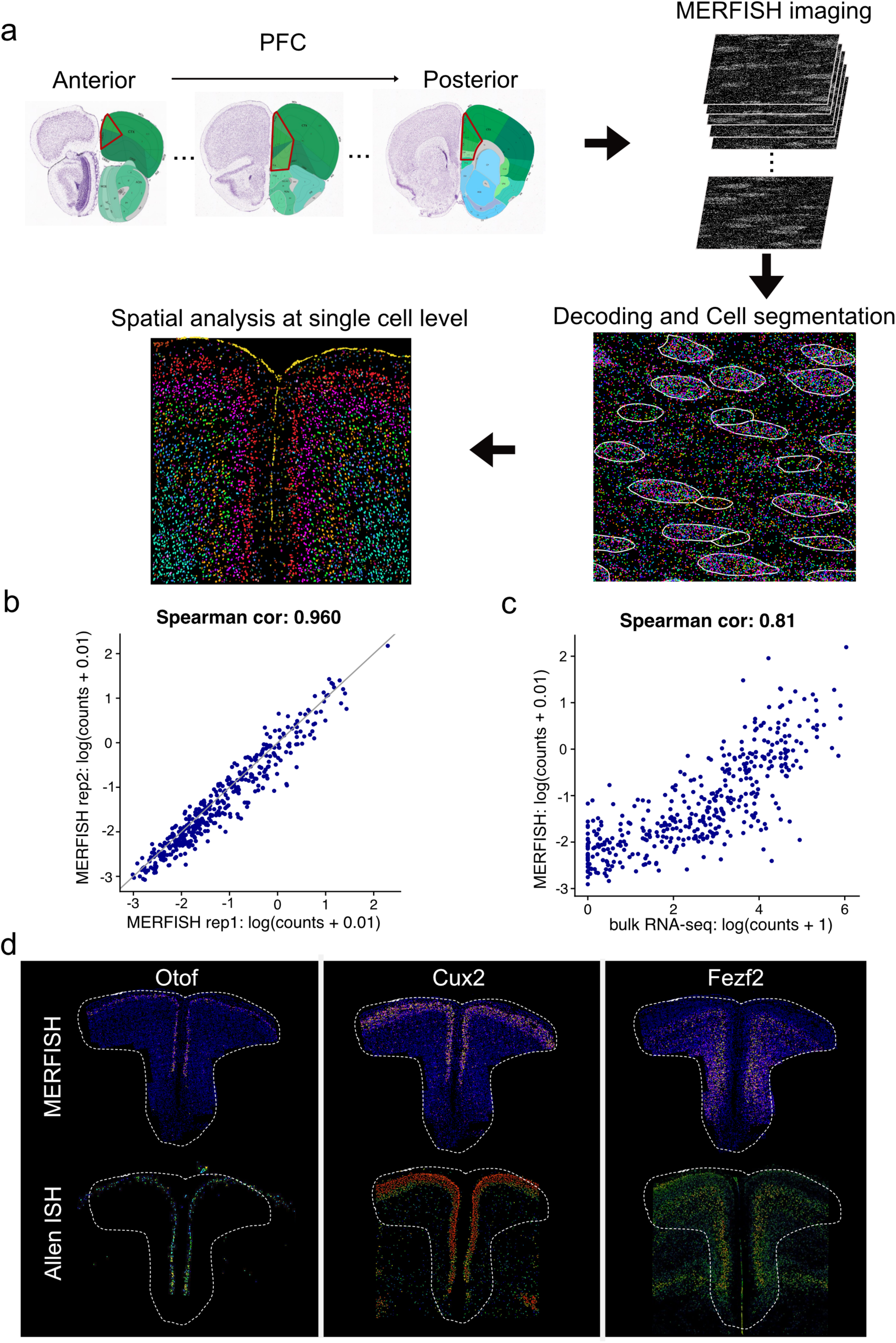
The workflow and quality control for MERFISH profiling. **a**, The workflow of MERFISH profiling of mouse PFC, including MERFISH imaging, decoding, segmentation and data analysis. **b**, Scatterplot showing the spearman correlation of the RNA counts per cell of individual genes measured by MERFISH in two independent experiments. **c**, Scatterplot of the RNA counts per cell of individual genes measured by MERFISH versus bulk RNA-seq data. The counts are natural logarithms. **d**, Spatial gene expression of three representative genes detected by MERFISH. *In situ* hybridization (ISH) data from Allen Brain Atlas are shown at the bottom.

**Fig. S2:**
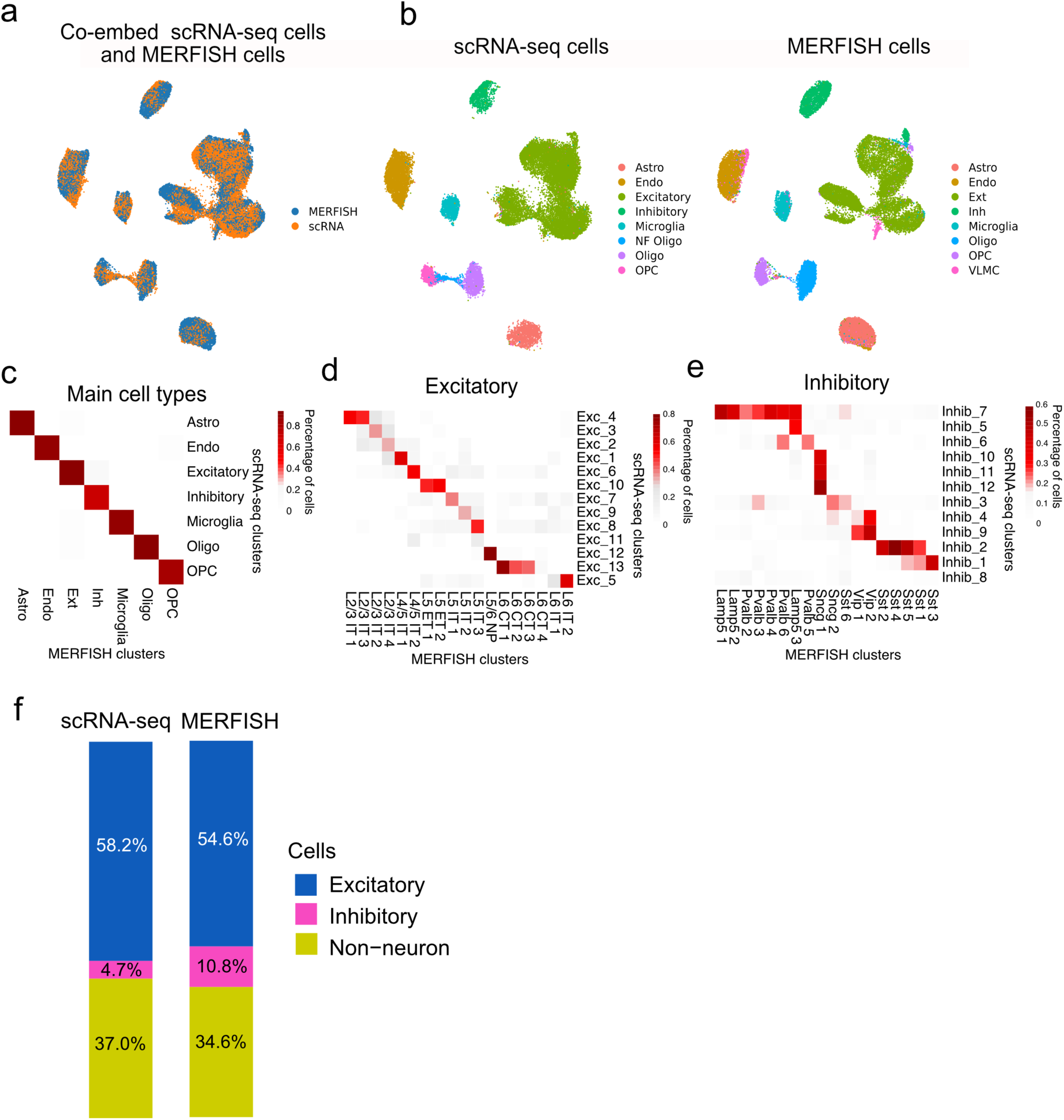
MERFISH and scRNA-seq based clusters are consistent. **a**, UMAP showing integration of cells from MERFISH or scRNA-seq data (GSE124952). **b**, UMAP showing the cell clusters defined by scRNA-seq (left) or MERFISH (right). **c**,**d**,**e**, Heatmap showing the correspondence between main cell types (**c**), excitatory (**d**) and inhibitory (**e**) subtypes defined by MERFISH and scRNA-seq. **f**, The cell proportions of the excitatory, inhibitory and non-neuronal cells from scRNA-seq or MERFISH.

**Fig. S3:**
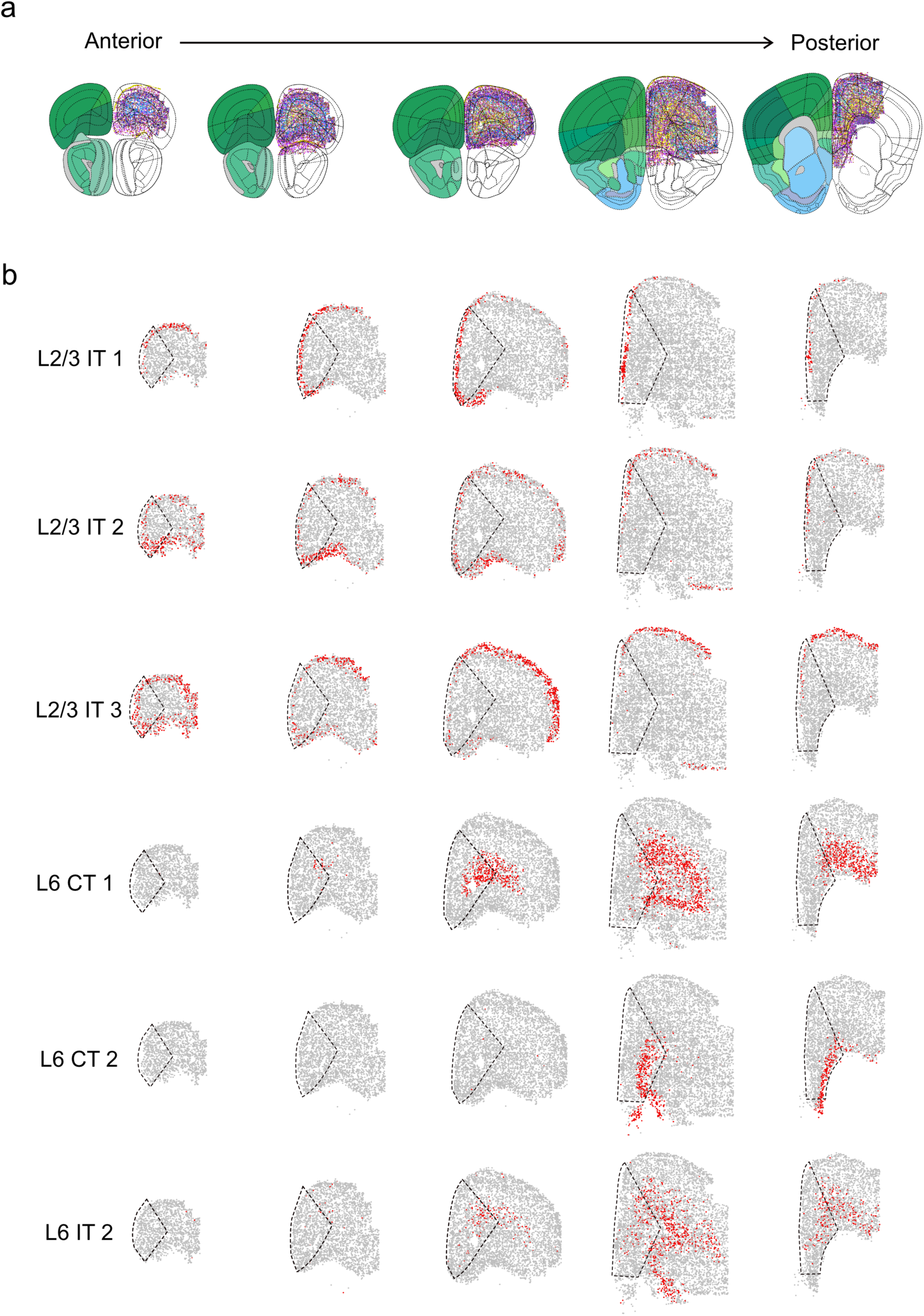
Spatial distribution of molecularly defined excitatory neuron subtypes along the anterior to posterior axis. **a**, Schematics of coronal brain slices aligned to Allen Brain Atlas CCF-v3 from anterior to posterior sections. **b**, Spatial organization of the indicated representative excitatory neuron subtypes across anterior to posterior sections.

**Fig. S4:**
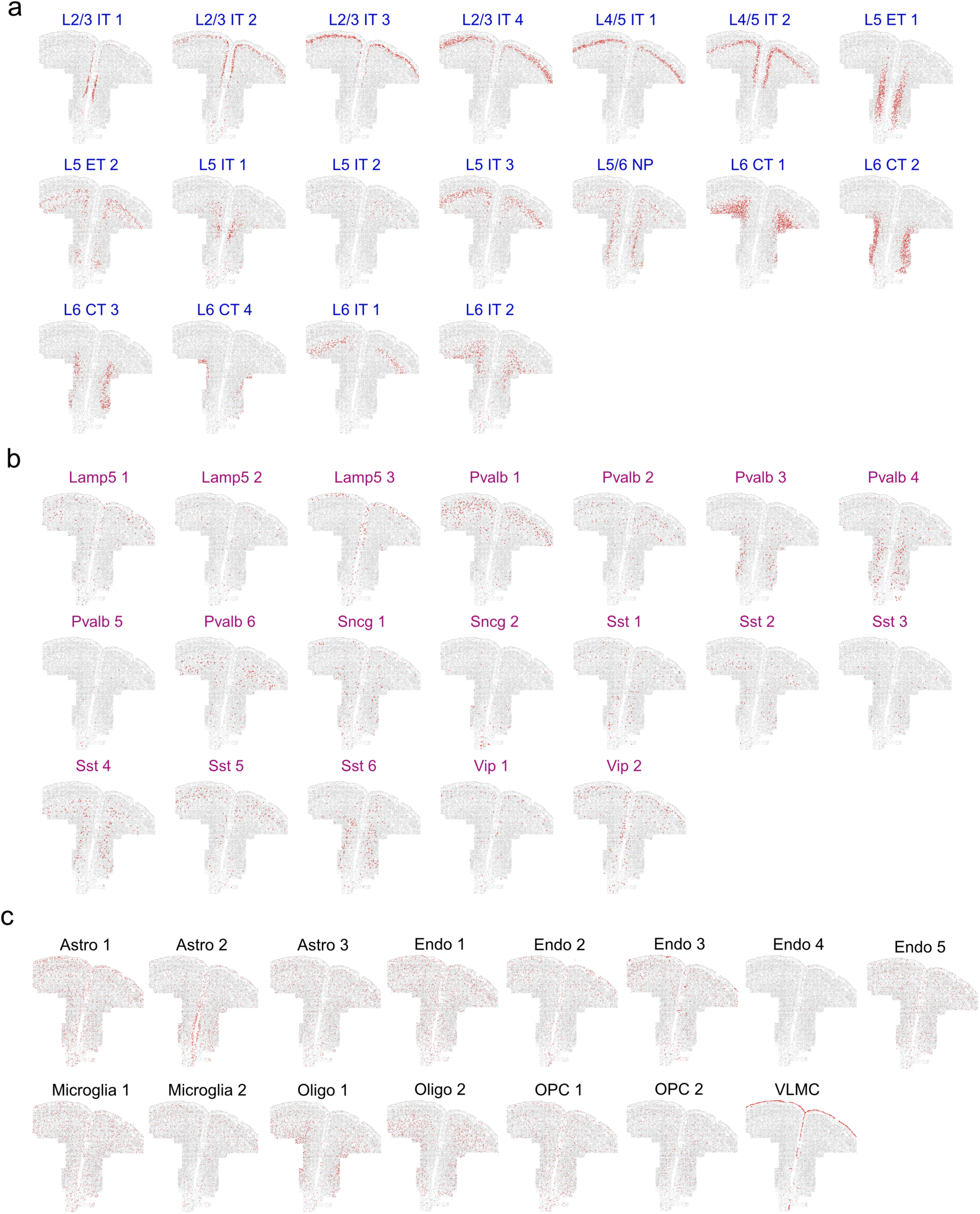
Spatial location of all molecularly defined PFC cell types and subtypes. **a**, Excitatory neuron subtypes; **b**, Inhibitory neuron subtypes; **c**, non-neuron cell types and subtypes. Red dots represent the indicated cell types and subtypes.

**Fig. S5:**
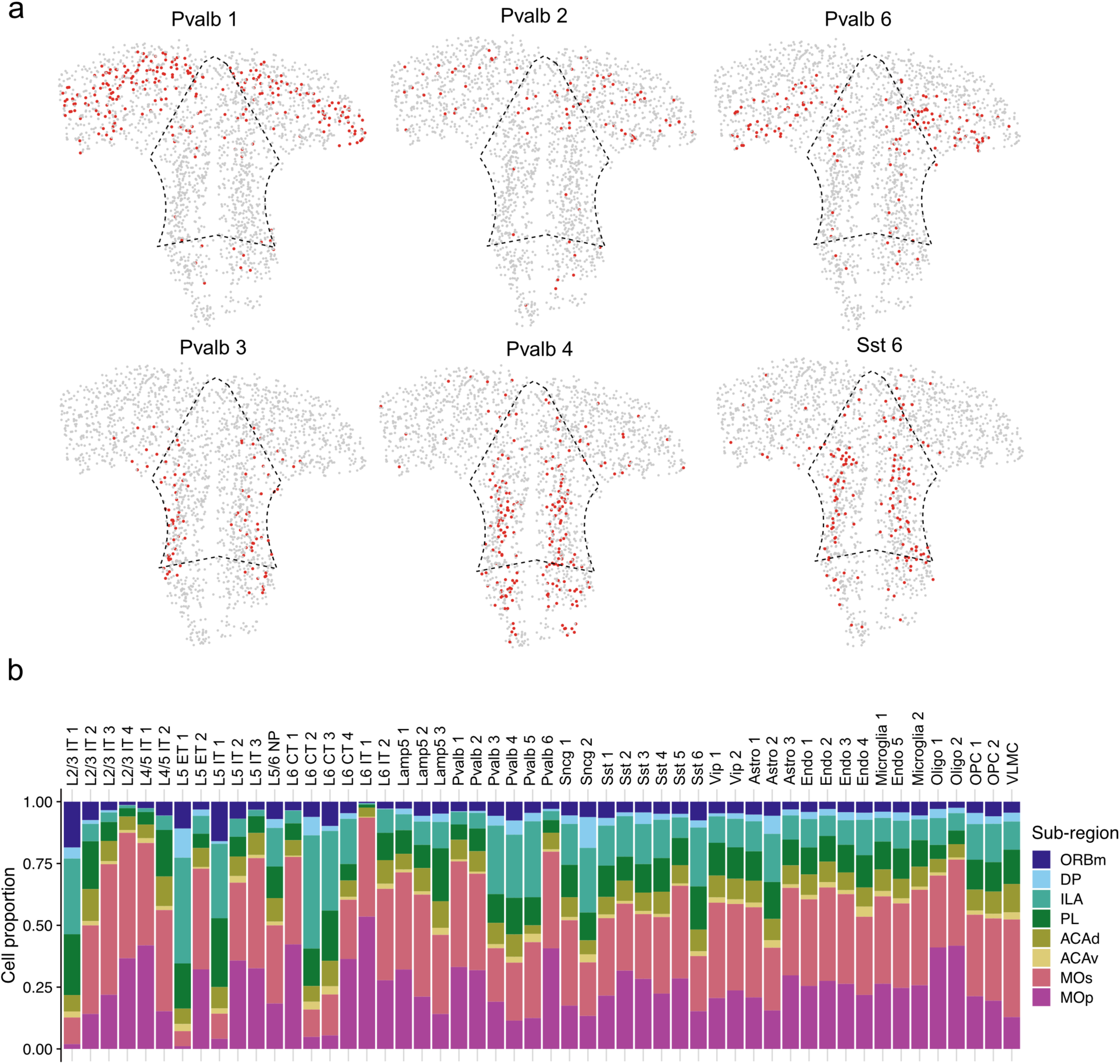
Distinct neuron subtypes are uniquely enriched in PFC and PFC subregions. **a,** Spatial location of three enriched (top panels: Pvalb 1, Pvalb 2, and Pvalb 6) and three depleted (bottom panels: Pvalb 3, Pvalb 4, and Sst 6) inhibitory subtypes on a coronal slice. **b**, The proportion of cell numbers from different PFC subregions and adjacent cortical regions of all neuron and non-neuron subtypes.

**Fig. S6:**
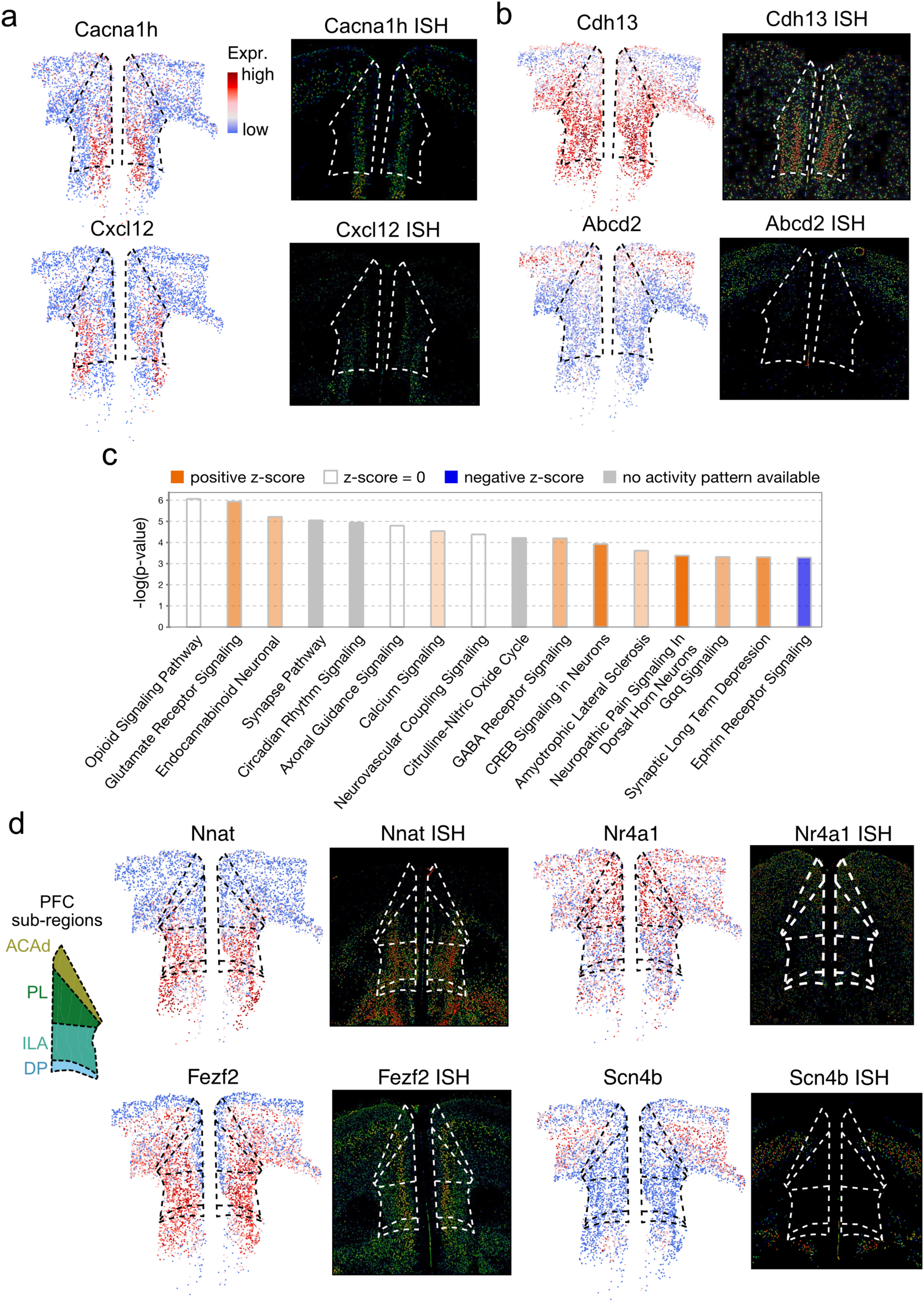
Specific gene expression signatures of PFC and PFC subregions. **a**,**b**, Spatial expression of two representative genes enriched (**a**) and depleted (**b**) in PFC relative to adjacent cortical regions. Only excitatory neurons are shown. Corresponding ISH data from Allen Brain Atlas are shown on the right. Dotted line marks PFC region. **c**, Ingenuity pathway analysis (IPA) of the genes, identified after imputation, showing enriched or depleted in PFC. The red/blue bars indicate the pathway more active in/out PFC, respectively. **d**, Spatial gene expression of four representative genes enriched in PFC subregions. A diagram of anatomical subregions in PFC and adjacent regions is shown on the left. Only the excitatory neurons are shown. ISH data from Allen Brain Atlas are shown on the right. Dotted line marks PFC subregion.

**Fig. S7:**
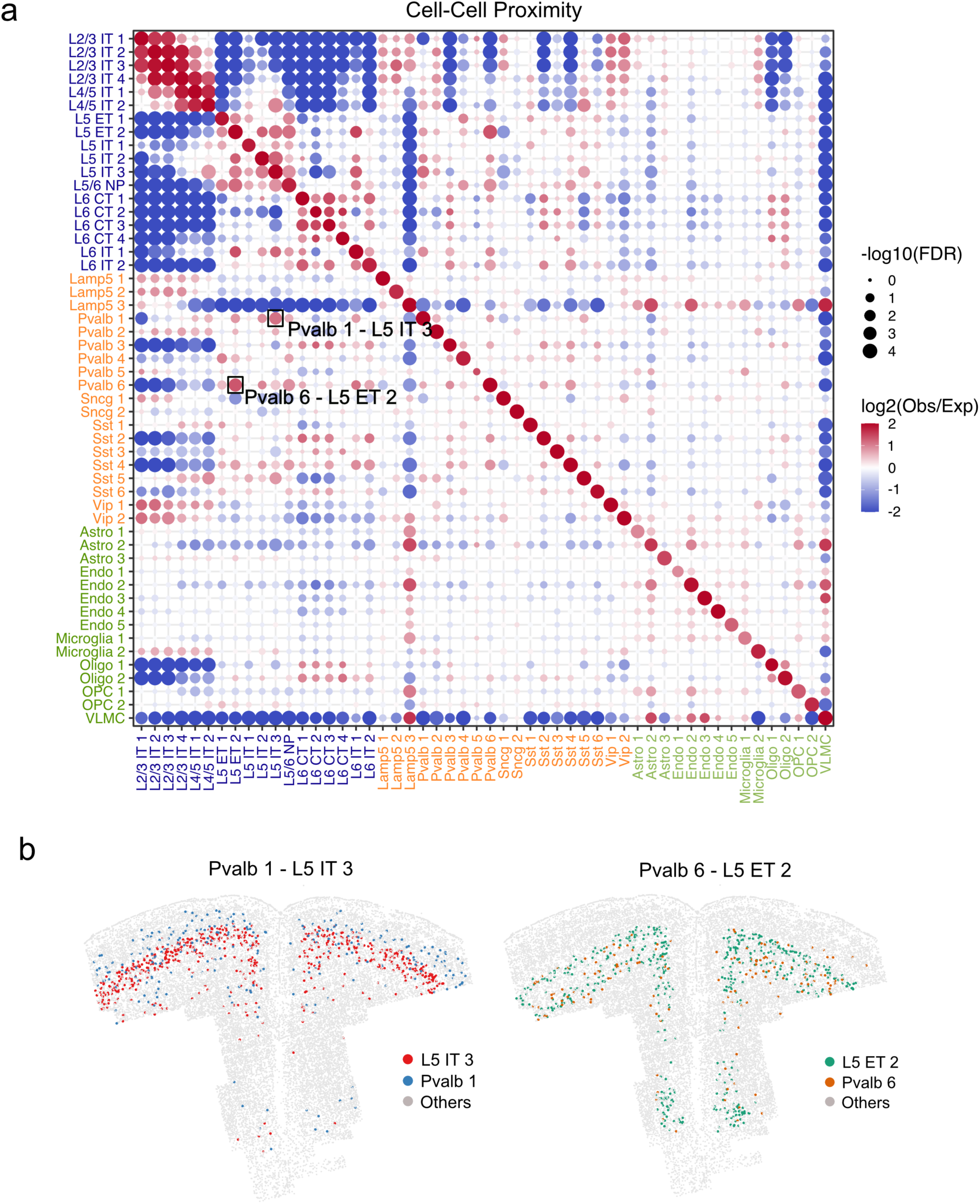
Cell-Cell proximity across all cell types. **a**, Enrichment of cell-cell proximity between different cell types and subtypes shown in dot plot. The color represents log2 transformed observation to expectation of co-localized frequency of two clusters. The size of dots indicates the significance of the co-localization. **b**, The cell-cell proximity between Pvalb 1 and L5 IT 3 neurons (left), and between Pvalb 6 and L5 ET 2 neurons (right).

**Fig. S8:**
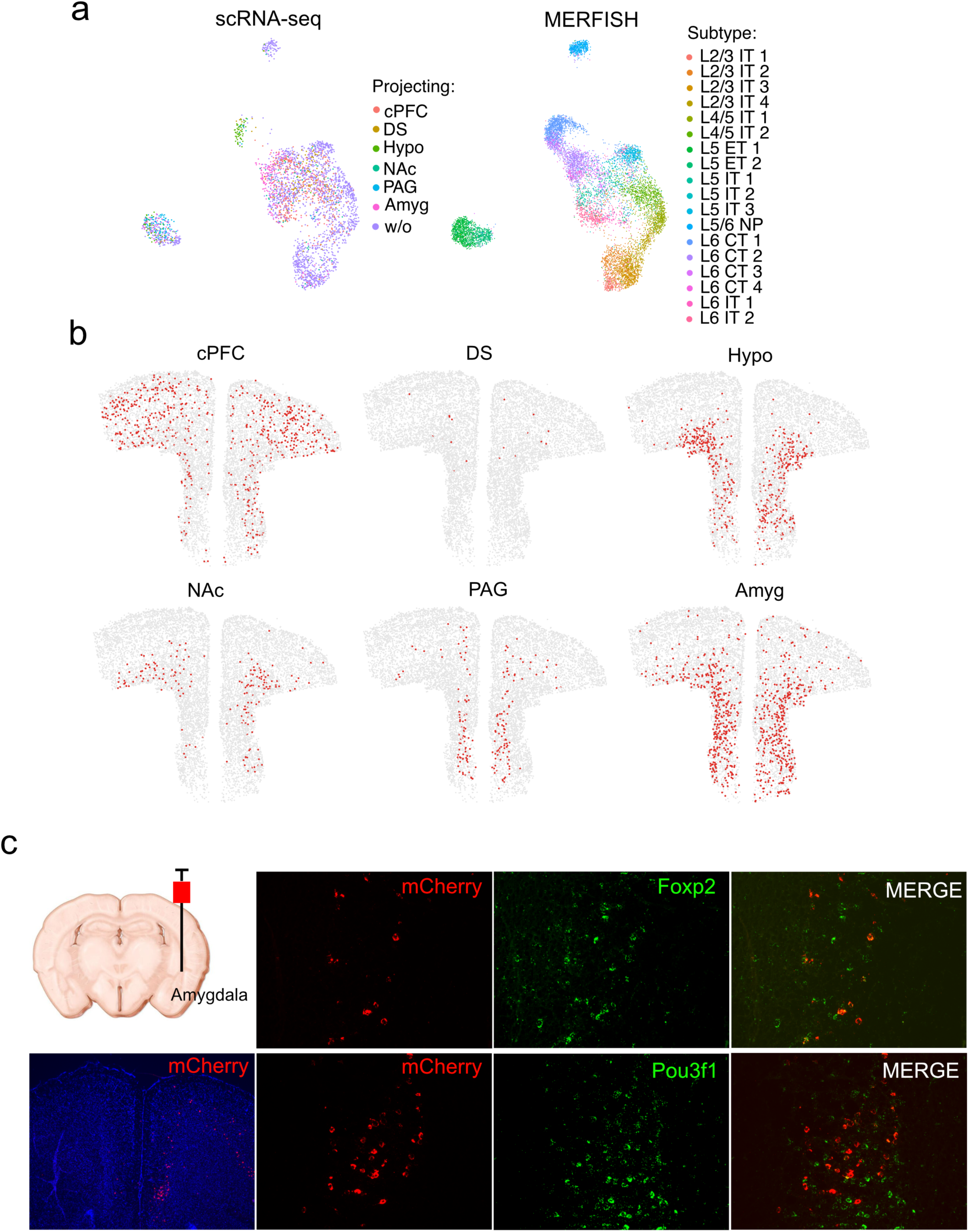
Integrate MERFISH and scRNA-seq data to predict neuronal projections. **a**, UMAP showing integration of cells from scRNA-seq (left) and MERFISH (right). The colors represent the projection sites in scRNA-seq data and the excitatory subtype in MERFISH data, respectively. **b**, Spatial location of neurons projecting to six different brain regions. **c**, Amygdala projection validation: mCherry expressing retrograde AAV was injected in amygdala. Brain slice of PFC were stained with DAPI and mCherry to image the labeled neurons. smFISH co-labeling of mCherry with *Pou3f1* (L5 ET marker), or *Foxp2* (L6 CT marker) reveal partial overlap with both neuron subtypes.

## List of Supplemental Tables

**Table S1: List of MERFISH probes**

**Table S2: List of enriched and depleted genes in PFC compared to the adjacent cortical regions**

**Table S3: List of genes whose expression is affected by chronic pain**

## References

1. Miller, E. K. & Cohen, J. D. An integrative theory of prefrontal cortex function. Annu Rev Neurosci 24, 167–202 (2001). https://doi.org:10.1146/annurev.neuro.24.1.167

2. Lui, J. H. et al. Differential encoding in prefrontal cortex projection neuron classes across cognitive tasks. Cell 184, 489–506 e426 (2021). https://doi.org:10.1016/j.cell.2020.11.046

3. Gamo, N. J. & Arnsten, A. F. Molecular modulation of prefrontal cortex: rational development of treatments for psychiatric disorders. Behav Neurosci 125, 282–296 (2011). https://doi.org:10.1037/a0023165

4. Chini, M. & Hanganu-Opatz, I. L. Prefrontal Cortex Development in Health and Disease: Lessons from Rodents and Humans. Trends Neurosci 44, 227–240 (2021). https://doi.org:10.1016/j.tins.2020.10.017

5. Tan, L. L. & Kuner, R. Neocortical circuits in pain and pain relief. Nat Rev Neurosci 22, 458–471 (2021). https://doi.org:10.1038/s41583-021-00468-2

6. Bushnell, M. C., Ceko, M. & Low, L. A. Cognitive and emotional control of pain and its disruption in chronic pain. Nat Rev Neurosci 14, 502–511 (2013). https://doi.org:10.1038/nrn3516

7. Yong, R. J., Mullins, P. M. & Bhattacharyya, N. Prevalence of chronic pain among adults in the United States. Pain 163, e328–e332 (2022). https://doi.org:10.1097/j.pain.0000000000002291

8. Gaskin, D. J. & Richard, P. The economic costs of pain in the United States. J Pain 13, 715–724 (2012). https://doi.org:10.1016/j.jpain.2012.03.009

9. Zhou, H. et al. A novel neuromodulation strategy to enhance the prefrontal control to treat pain. Mol Pain 15, 1744806919845739 (2019). https://doi.org:10.1177/1744806919845739

10. Deer, T. R. et al. The appropriate use of neurostimulation: new and evolving neurostimulation therapies and applicable treatment for chronic pain and selected disease states. Neuromodulation Appropriateness Consensus Committee. Neuromodulation 17, 599–615; discussion 615 (2014). https://doi.org:10.1111/ner.12204

11. Baliki, M. N., Geha, P. Y., Apkarian, A. V. & Chialvo, D. R. Beyond feeling: chronic pain hurts the brain, disrupting the default-mode network dynamics. J Neurosci 28, 1398–1403 (2008). https://doi.org:10.1523/JNEUROSCI.4123-07.2008

12. Davis, K. D. & Moayedi, M. Central mechanisms of pain revealed through functional and structural MRI. J Neuroimmune Pharmacol 8, 518–534 (2013). https://doi.org:10.1007/s11481-012-9386-8

13. Nardone, R. et al. rTMS of the prefrontal cortex has analgesic effects on neuropathic pain in subjects with spinal cord injury. Spinal Cord 55, 20–25 (2017). https://doi.org:10.1038/sc.2016.87

14. Jefferson, T., Kelly, C. J. & Martina, M. Differential Rearrangement of Excitatory Inputs to the Medial Prefrontal Cortex in Chronic Pain Models. Front Neural Circuits 15, 791043 (2021). https://doi.org:10.3389/fncir.2021.791043

15. Li, C. et al. Prolonged Continuous Theta Burst Stimulation Can Regulate Sensitivity on Abeta Fibers: An Functional Near-Infrared Spectroscopy Study. Front Mol Neurosci 15, 887426 (2022). https://doi.org:10.3389/fnmol.2022.887426

16. Ong, W. Y., Stohler, C. S. & Herr, D. R. Role of the Prefrontal Cortex in Pain Processing. Mol Neurobiol 56, 1137–1166 (2019). https://doi.org:10.1007/s12035-018-1130-9

17. Ossipov, M. H., Morimura, K. & Porreca, F. Descending pain modulation and chronification of pain. Curr Opin Support Palliat Care 8, 143–151 (2014). https://doi.org:10.1097/SPC.0000000000000055

18. Anastasiades, P. G. & Carter, A. G. Circuit organization of the rodent medial prefrontal cortex. Trends Neurosci 44, 550–563 (2021). https://doi.org:10.1016/j.tins.2021.03.006

19. Bhattacherjee, A. et al. Cell type-specific transcriptional programs in mouse prefrontal cortex during adolescence and addiction. Nat Commun 10, 4169 (2019). https://doi.org:10.1038/s41467-019-12054-3

20. Zeng, H. & Sanes, J. R. Neuronal cell-type classification: challenges, opportunities and the path forward. Nat Rev Neurosci 18, 530–546 (2017). https://doi.org:10.1038/nrn.2017.85

21. Network, B. I. C. C. A multimodal cell census and atlas of the mammalian primary motor cortex. Nature 598, 86–102 (2021). https://doi.org:10.1038/s41586-021-03950-0

22. Radnikow, G. & Feldmeyer, D. Layer- and Cell Type-Specific Modulation of Excitatory Neuronal Activity in the Neocortex. Front Neuroanat 12, 1 (2018). https://doi.org:10.3389/fnana.2018.00001

23. Tasic, B. et al. Shared and distinct transcriptomic cell types across neocortical areas. Nature 563, 72–78 (2018). https://doi.org:10.1038/s41586-018-0654-5

24. Loo, L. et al. Single-cell transcriptomic analysis of mouse neocortical development. Nat Commun 10, 134 (2019). https://doi.org:10.1038/s41467-018-08079-9

25. Li, Y. E. et al. An atlas of gene regulatory elements in adult mouse cerebrum. Nature 598, 129–136 (2021). https://doi.org:10.1038/s41586-021-03604-1

26. Chen, K. H., Boettiger, A. N., Moffitt, J. R., Wang, S. & Zhuang, X. RNA imaging. Spatially resolved, highly multiplexed RNA profiling in single cells. Science 348, aaa6090 (2015). https://doi.org:10.1126/science.aaa6090

27. Xia, C., Fan, J., Emanuel, G., Hao, J. & Zhuang, X. Spatial transcriptome profiling by MERFISH reveals subcellular RNA compartmentalization and cell cycle-dependent gene expression. Proc Natl Acad Sci U S A 116, 19490–19499 (2019). https://doi.org:10.1073/pnas.1912459116

28. Moffitt, J. R. & Zhuang, X. RNA Imaging with Multiplexed Error-Robust Fluorescence In Situ Hybridization (MERFISH). Methods Enzymol 572, 1–49 (2016). https://doi.org:10.1016/bs.mie.2016.03.020

29. Moffitt, J. R. et al. Molecular, spatial, and functional single-cell profiling of the hypothalamic preoptic region. Science 362 (2018). https://doi.org:10.1126/science.aau5324

30. Wang, Q. et al. The Allen Mouse Brain Common Coordinate Framework: A 3D Reference Atlas. Cell 181, 936–953 e920 (2020). https://doi.org:10.1016/j.cell.2020.04.007

31. Oishi, K. et al. Identity of neocortical layer 4 neurons is specified through correct positioning into the cortex. Elife 5 (2016). https://doi.org:10.7554/eLife.10907

32. Zhang, M. et al. Spatially resolved cell atlas of the mouse primary motor cortex by MERFISH. Nature 598, 137–143 (2021). https://doi.org:10.1038/s41586-021-03705-x

33. Isaacson, J. S. & Scanziani, M. How inhibition shapes cortical activity. Neuron 72, 231–243 (2011). https://doi.org:10.1016/j.neuron.2011.09.027

34. Jang, H. J. et al. Distinct roles of parvalbumin and somatostatin interneurons in gating the synchronization of spike times in the neocortex. Sci Adv 6, eaay5333 (2020). https://doi.org:10.1126/sciadv.aay5333

35. Brown, J. A. et al. Inhibition of parvalbumin-expressing interneurons results in complex behavioral changes. Mol Psychiatry 20, 1499–1507 (2015). https://doi.org:10.1038/mp.2014.192

36. Santini, E., Quirk, G. J. & Porter, J. T. Fear conditioning and extinction differentially modify the intrinsic excitability of infralimbic neurons. J Neurosci 28, 4028–4036 (2008). https://doi.org:10.1523/JNEUROSCI.2623-07.2008

37. Otis, J. M. et al. Prefrontal cortex output circuits guide reward seeking through divergent cue encoding. Nature 543, 103–107 (2017). https://doi.org:10.1038/nature21376

38. Spring, M. G., Soni, K. R., Wheeler, D. S. & Wheeler, R. A. Prelimbic prefrontal cortical encoding of reward predictive cues. Synapse 75, e22202 (2021). https://doi.org:10.1002/syn.22202

39. Harada, M., Pascoli, V., Hiver, A., Flakowski, J. & Luscher, C. Corticostriatal Activity Driving Compulsive Reward Seeking. Biol Psychiatry 90, 808–818 (2021). https://doi.org:10.1016/j.biopsych.2021.08.018

40. Humphries, E. S. & Dart, C. Neuronal and Cardiovascular Potassium Channels as Therapeutic Drug Targets: Promise and Pitfalls. J Biomol Screen 20, 1055–1073 (2015). https://doi.org:10.1177/1087057115601677

41. Gonzalez Sabater, V., Rigby, M. & Burrone, J. Voltage-Gated Potassium Channels Ensure Action Potential Shape Fidelity in Distal Axons. J Neurosci 41, 5372–5385 (2021). https://doi.org:10.1523/JNEUROSCI.2765-20.2021

42. Lai, H. C. & Jan, L. Y. The distribution and targeting of neuronal voltage-gated ion channels. Nat Rev Neurosci 7, 548–562 (2006). https://doi.org:10.1038/nrn1938

43. Grube, S. et al. A CAG repeat polymorphism of KCNN3 predicts SK3 channel function and cognitive performance in schizophrenia. EMBO Mol Med 3, 309–319 (2011). https://doi.org:10.1002/emmm.201100135

44. Andrade, A. et al. Genetic Associations between Voltage-Gated Calcium Channels and Psychiatric Disorders. Int J Mol Sci 20 (2019). https://doi.org:10.3390/ijms20143537

45. Eckle, V. S. et al. Mechanisms by which a CACNA1H mutation in epilepsy patients increases seizure susceptibility. J Physiol 592, 795–809 (2014). https://doi.org:10.1113/jphysiol.2013.264176

46. Carvill, G. L. Calcium Channel Dysfunction in Epilepsy: Gain of CACNA1E. Epilepsy Curr 19, 199–201 (2019). https://doi.org:10.1177/1535759719845324

47. Splawski, I. et al. CACNA1H mutations in autism spectrum disorders. J Biol Chem 281, 22085–22091 (2006). https://doi.org:10.1074/jbc.M603316200

48. Zhang, J. & Abdullah, J. M. The role of GluA1 in central nervous system disorders. Rev Neurosci 24, 499–505 (2013). https://doi.org:10.1515/revneuro-2013-0021

49. Qu, W. et al. Emerging role of AMPA receptor subunit GluA1 in synaptic plasticity: Implications for Alzheimer’s disease. Cell Prolif 54, e12959 (2021). https://doi.org:10.1111/cpr.12959

50. Forrest, M. P., Parnell, E. & Penzes, P. Dendritic structural plasticity and neuropsychiatric disease. Nat Rev Neurosci 19, 215–234 (2018). https://doi.org:10.1038/nrn.2018.16

51. Peng, S. X. et al. SNP rs10420324 in the AMPA receptor auxiliary subunit TARP gamma-8 regulates the susceptibility to antisocial personality disorder. Sci Rep 11, 11997 (2021). https://doi.org:10.1038/s41598-021-91415-9

52. Festa, L. K. et al. CXCL12-induced rescue of cortical dendritic spines and cognitive flexibility. Elife 9 (2020). https://doi.org:10.7554/eLife.49717

53. Wu, P. R., Cho, K. K. A., Vogt, D., Sohal, V. S. & Rubenstein, J. L. R. The Cytokine CXCL12 Promotes Basket Interneuron Inhibitory Synapses in the Medial Prefrontal Cortex. Cereb Cortex 27, 4303–4313 (2017). https://doi.org:10.1093/cercor/bhw230

54. Sanfilippo, C., Castrogiovanni, P., Imbesi, R., Nunnari, G. & Di Rosa, M. Postsynaptic damage and microglial activation in AD patients could be linked CXCR4/CXCL12 expression levels. Brain Res 1749, 147127 (2020). https://doi.org:10.1016/j.brainres.2020.147127

55. Zhang, C., Chen, R. & Zhang, Y. Accurate inference of genome-wide spatial expression with iSpatial. Sci Adv 8, eabq0990 (2022). https://doi.org:10.1126/sciadv.abq0990

56. Kurowski, P., Grzelka, K. & Szulczyk, P. Ionic Mechanism Underlying Rebound Depolarization in Medial Prefrontal Cortex Pyramidal Neurons. Front Cell Neurosci 12, 93 (2018). https://doi.org:10.3389/fncel.2018.00093

57. Selleck, R. A. et al. Endogenous Opioid Signaling in the Medial Prefrontal Cortex is Required for the Expression of Hunger-Induced Impulsive Action. Neuropsychopharmacology 40, 2464–2474 (2015). https://doi.org:10.1038/npp.2015.97

58. Baldo, B. A. Prefrontal Cortical Opioids and Dysregulated Motivation: A Network Hypothesis. Trends Neurosci 39, 366–377 (2016). https://doi.org:10.1016/j.tins.2016.03.004

59. Tan, H., Ahmad, T., Loureiro, M., Zunder, J. & Laviolette, S. R. The role of cannabinoid transmission in emotional memory formation: implications for addiction and schizophrenia. Front Psychiatry 5, 73 (2014). https://doi.org:10.3389/fpsyt.2014.00073

60. Egerton, A., Allison, C., Brett, R. R. & Pratt, J. A. Cannabinoids and prefrontal cortical function: insights from preclinical studies. Neurosci Biobehav Rev 30, 680–695 (2006). https://doi.org:10.1016/j.neubiorev.2005.12.002

61. Peron, S. et al. Recurrent interactions in local cortical circuits. Nature 579, 256–259 (2020). https://doi.org:10.1038/s41586-020-2062-x

62. LaBerge, D. & Kasevich, R. S. Neuroelectric Tuning of Cortical Oscillations by Apical Dendrites in Loop Circuits. Front Syst Neurosci 11, 37 (2017). https://doi.org:10.3389/fnsys.2017.00037

63. Lacefield, C. O., Pnevmatikakis, E. A., Paninski, L. & Bruno, R. M. Reinforcement Learning Recruits Somata and Apical Dendrites across Layers of Primary Sensory Cortex. Cell Rep 26, 2000–2008 e2002 (2019). https://doi.org:10.1016/j.celrep.2019.01.093

64. Karimi, A., Odenthal, J., Drawitsch, F., Boergens, K. M. & Helmstaedter, M. Cell-type specific innervation of cortical pyramidal cells at their apical dendrites. Elife 9 (2020). https://doi.org:10.7554/eLife.46876

65. Harris, K. D. & Shepherd, G. M. The neocortical circuit: themes and variations. Nat Neurosci 18, 170–181 (2015). https://doi.org:10.1038/nn.3917

66. Wamsley, B. & Fishell, G. Genetic and activity-dependent mechanisms underlying interneuron diversity. Nat Rev Neurosci 18, 299–309 (2017). https://doi.org:10.1038/nrn.2017.30

67. Gabbott, P. L., Warner, T. A., Jays, P. R., Salway, P. & Busby, S. J. Prefrontal cortex in the rat: projections to subcortical autonomic, motor, and limbic centers. J Comp Neurol 492, 145–177 (2005). https://doi.org:10.1002/cne.20738

68. Zingg, B. et al. Neural networks of the mouse neocortex. Cell 156, 1096–1111 (2014). https://doi.org:10.1016/j.cell.2014.02.023

69. Baxter, M. G. & Croxson, P. L. Facing the role of the amygdala in emotional information processing. Proc Natl Acad Sci U S A 109, 21180–21181 (2012). https://doi.org:10.1073/pnas.1219167110

70. Bonnet, L. et al. The role of the amygdala in the perception of positive emotions: an “intensity detector”. Front Behav Neurosci 9, 178 (2015). https://doi.org:10.3389/fnbeh.2015.00178

71. Corder, G. et al. An amygdalar neural ensemble that encodes the unpleasantness of pain. Science 363, 276–281 (2019). https://doi.org:10.1126/science.aap8586

72. Topham, L. et al. The transition from acute to chronic pain: dynamic epigenetic reprogramming of the mouse prefrontal cortex up to 1 year after nerve injury. Pain 161, 2394–2409 (2020). https://doi.org:10.1097/j.pain.0000000000001917

73. Descalzi, G. et al. Neuropathic pain promotes adaptive changes in gene expression in brain networks involved in stress and depression. Sci Signal 10 (2017). https://doi.org:10.1126/scisignal.aaj1549

74. Richner, M., Bjerrum, O. J., Nykjaer, A. & Vaegter, C. B. The spared nerve injury (SNI) model of induced mechanical allodynia in mice. J Vis Exp (2011). https://doi.org:10.3791/3092

75. Denk, F., McMahon, S. B. & Tracey, I. Pain vulnerability: a neurobiological perspective. Nat Neurosci 17, 192–200 (2014). https://doi.org:10.1038/nn.3628

76. Moffitt, J. R., Lundberg, E. & Heyn, H. The emerging landscape of spatial profiling technologies. Nat Rev Genet 23, 741–759 (2022). https://doi.org:10.1038/s41576-022-00515-3

77. Chen, R. et al. Decoding molecular and cellular heterogeneity of mouse nucleus accumbens. Nat Neurosci 24, 1757–1771 (2021). https://doi.org:10.1038/s41593-021-00938-x

78. Collins, D. P., Anastasiades, P. G., Marlin, J. J. & Carter, A. G. Reciprocal Circuits Linking the Prefrontal Cortex with Dorsal and Ventral Thalamic Nuclei. Neuron 98, 366–379 e364 (2018). https://doi.org:10.1016/j.neuron.2018.03.024

79. Song, C. & Moyer, J. R., Jr. Layer- and subregion-specific differences in the neurophysiological properties of rat medial prefrontal cortex pyramidal neurons. J Neurophysiol 119, 177–191 (2018). https://doi.org:10.1152/jn.00146.2017

80. Lodge, D. J., Behrens, M. M. & Grace, A. A. A loss of parvalbumin-containing interneurons is associated with diminished oscillatory activity in an animal model of schizophrenia. J Neurosci 29, 2344–2354 (2009). https://doi.org:10.1523/JNEUROSCI.5419-08.2009

81. Zhang, Z. et al. Role of Prelimbic GABAergic Circuits in Sensory and Emotional Aspects of Neuropathic Pain. Cell Rep 12, 752–759 (2015). https://doi.org:10.1016/j.celrep.2015.07.001

82. Gallego-Carracedo, C., Perich, M. G., Chowdhury, R. H., Miller, L. E. & Gallego, J. A. Local field potentials reflect cortical population dynamics in a region-specific and frequency-dependent manner. Elife 11 (2022). https://doi.org:10.7554/eLife.73155

83. Herreras, O. Local Field Potentials: Myths and Misunderstandings. Front Neural Circuits 10, 101 (2016). https://doi.org:10.3389/fncir.2016.00101

84. Liu, Y. et al. Frequency Dependent Electrical Stimulation of PFC and ACC for Acute Pain Treatment in Rats. Front Pain Res (Lausanne) 2, 728045 (2021). https://doi.org:10.3389/fpain.2021.728045

85. Dale, J. et al. Scaling Up Cortical Control Inhibits Pain. Cell Rep 23, 1301–1313 (2018). https://doi.org:10.1016/j.celrep.2018.03.139

86. Baliki, M. N., Geha, P. Y., Fields, H. L. & Apkarian, A. V. Predicting value of pain and analgesia: nucleus accumbens response to noxious stimuli changes in the presence of chronic pain. Neuron 66, 149–160 (2010). https://doi.org:10.1016/j.neuron.2010.03.002

87. Generaal, E. et al. Reduced hypothalamic-pituitary-adrenal axis activity in chronic multi-site musculoskeletal pain: partly masked by depressive and anxiety disorders. BMC Musculoskelet Disord 15, 227 (2014). https://doi.org:10.1186/1471-2474-15-227

88. Wolock, S. L., Lopez, R. & Klein, A. M. Scrublet: Computational Identification of Cell Doublets in Single-Cell Transcriptomic Data. Cell Syst 8, 281–291 e289 (2019). https://doi.org:10.1016/j.cels.2018.11.005

89. Hao, Y. et al. Integrated analysis of multimodal single-cell data. Cell 184, 3573–3587 e3529 (2021). https://doi.org:10.1016/j.cell.2021.04.048

90. Korsunsky, I. et al. Fast, sensitive and accurate integration of single-cell data with Harmony. Nat Methods 16, 1289–1296 (2019). https://doi.org:10.1038/s41592-019-0619-0

91. Traag, V. A., Waltman, L. & van Eck, N. J. From Louvain to Leiden: guaranteeing well-connected communities. Sci Rep 9, 5233 (2019). https://doi.org:10.1038/s41598-019-41695-z

92. Furth, D. et al. An interactive framework for whole-brain maps at cellular resolution. Nat Neurosci 21, 139–149 (2018). https://doi.org:10.1038/s41593-017-0027-7

